# Particle Phagocytosis Amplifies Secretion of Small Extracellular Vesicles in Macrophages

**DOI:** 10.64898/2026.01.29.701815

**Authors:** Michael Trautmann-Rodriguez, Vilina Akala, Nicole A. Gill, Emma R. Sudduth, Catherine A. Fromen

**Affiliations:** Department of Chemical and Biomolecular Engineering, University of Delaware, Newark, DE 19716; Department of Biomedical Engineering, University of Delaware, Newark, DE 19716

## Abstract

Macrophage-derived extracellular vesicles (EVs) are critical mediators of immune communication and tissue homeostasis, yet the exogenous factors that regulate their release and composition remain poorly defined. Here, we identify phagocytosis of particulate materials as a direct and general driver of macrophage small EV production through a lysosome-linked mechanism. Across primary and engineered murine and human macrophage cultures, phagocytic uptake robustly increased small EV secretion, an effect that was recapitulated *in vivo* following orotracheal instillation that resulted in elevated small EV abundance in bronchoalveolar lavage fluid. This phagocytosis-driven small EV amplification depended on particle size (hydrodynamic diameter >200 nm) and residence time within the phagolysosome. Electron microscopy and proteomic analyses revealed increased multivesicular body formation following particle uptake, establishing a mechanistic link between phagocytosis and small EV biogenesis. Notably, this pathway regulated both protein and particulate cargo in secreted macrophage small EVs. Together, these findings expand the canonical framework of macrophage EV biogenesis to include phagocytosis-driven regulation and establish a single step process for controlling small EV production and cargo loading, with implications for understanding particulate exposures and for developing macrophage- and EV-based immunotherapies.

## INTRODUCTION

Extracellular vesicles (EVs) are membrane-bound carriers that transport bioactive proteins, lipids, and nucleic acids between cells^1, 2^ and serve as essential mediators of immune communication.^2^ Current consensus categorizes EVs based on size, including small EVs (<200 nm) and large EVs (>200 nm).^1^ Small EVs are thought to originate from endosomal processing and multivesicular body (MVB) formation,^3, 4^ facilitating the transfer of critical RNA, DNA, and protein between cells in response to environmental cues.^1, 2^ Among immune cells, macrophages are prolific producers of EVs, with EV quantity and composition reflecting the state of the producing cell and mediating downstream immune signals.^5, 6^ Accordingly, changes in macrophage EV release and cargo can act as sensitive indicators of how these cells integrate environmental cues and respond to perturbations.^7–9^ Consistent with this, dysfunctional macrophage small EVs are increasingly implicated in various diseases, where altered composition and cargo transfer components contribute to pathogenesis.^7, 8^ Yet, the upstream mechanisms that drive the secretion of such dysfunctional macrophage EVs remain poorly defined.

Macrophages are also the dominant cell type responsible for clearing particulate materials, including environmental and engineered therapeutic nanoparticles (NP).^10, 11^ Following internalization, phagocytosed particles are trafficked through the phagolysosome, a pathway widely regarded as the terminal intracellular fate for exogenous material.^12, 13^ This degradative process is central to macrophage function, enabling efficient clearance of foreign material. However, late endosomal and lysosomal compartments are also associated with intracellular antigen-processing compartments that are functional MVBs, though the spatial, temporal, and mechanistic relationships between these compartments remain unresolved.^14^ These antigen-processing MVB compartments support immune communication and are known to give rise to EV secretion.^9, 14–16^ Despite extensive molecular characterization of phagosome maturation, lysosomal trafficking, and MVB exocytosis involving protein families such as chaperone proteins, small GTPases, and SNARE proteins,^14, 17, 18^ the functional connection between phagosome processing of particulate material and EV loading and biogenesis remains unclear.

As phagocyte uptake is a near-universal fate of particles administered *in vivo*,^19^ understanding the full downstream consequences and relationship to EV biogenesis is essential.^14^ Existing studies of macrophage-particle interactions have largely focused on particle uptake, intracellular trafficking, and inflammatory signaling outcomes.^13, 20–22^ Prior observations suggest that increased lysosomal activity correlates with increased EV production,^23^ and a small number of reports document EV changes after particle exposure, particularly in environmental or toxicological studies.^24^ However, these studies typically focus on single materials, do not distinguish uptake pathways, and rarely assess whether specific particle properties govern the EV output.^25, 26^ Consequently, whether phagocytosis itself acts as a regulatory input to macrophage EV biogenesis, rather than a concurrent inflammatory stimulus, has not been systematically tested.

We hypothesized that the phagocytic uptake and lysosomal persistence of particulate matter act as primary drivers of EV biogenesis in macrophages. To test this hypothesis, we designed a particle library that systematically varied size, chemistry, charge, density, and degradability and evaluated EV production and composition, biogenesis pathways, and EV-mediated material transfer. Together, our results reveal a conserved mechanism in which phagocytic uptake and intracellular material accumulation dictate macrophage small EV secretion, establishing a mechanistic framework linking phagocytosis to small EV biogenesis and providing design principles for engineering EV-mediated communication and NP fate.

## RESULTS

### Negatively-charged PEGDA NPs drive secretion of macrophage EVs *in vitro* and *in vivo*

To investigate the role of particle dosing in macrophage EV output, we first generated biocompatible hydrogel NPs ∼250 nm in diameter comprised of poly(ethylene glycol) diacrylate (PEGDA) formulated at 50 wt% solids with an anionic comonomer [PEGDA(-)50] consistent with prior studies (**Table 1**).^27^ PEGDA(-)50 was administered to various macrophages *in vitro*, and the resultant EVs were isolated 72 hrs after dosing and quantified for size and concentration yield. As shown in **Figure 1A**, RAW 264.7 *in vitro* cultures demonstrated an increase in secreted EVs following dosing with 50 to 400 µg/mL of PEGDA(-)50. **Supplemental Figure S1.I** confirms PEGDA(-)50 internalization following 24 hrs in RAW 264.7 cells. Cell cultures dosed with 100 µg/mL of PEGDA(-)50 showed a 5-fold increase in EV concentration following isolation via ultracentrifugation, while the RAW 264.7 cell cultures that received the highest dose (400 µg/mL) showed approximately a 30-fold increase in EV concentration (p = 0.0145 and p<0.0001 respectively, via ordinary one-way ANOVA with Turkey’s multiple comparisons test). Highlighted as an example, EVs from RAW 264.7 macrophage cultures dosed with 100 µg/mL have a diameter mode and mean (91.4 ± 1.1 nm and 103.1 ± 1.1 nm, respectively) that is characteristic of small EVs. These results point to a direct relationship between RAW 264.7 macrophage EVs and the dose concentration of PEGDA(-)50.

**Table 1.** Particle Chemistry and Characteristics. The table includes the average hydrodynamic diameter (D_h_), polydispersity index (PDI), and overall expected charge for the particle formulations used throughout the study.

| Name | PEGDA (%) | dS (%) | +/- | wt% | D<sub>h</sub> (nm) | PDI |
| --- | --- | --- | --- | --- | --- | --- |
| PEGDA(-)50 [NP] | 88.8 | 0 | - | 50 | 240.9 ± 26.0 | 0.19 ± 0.09 |
| PEGDA(-)75 | 88.8 | 0 | - | 75 | 247.2 ± 0.35 | 0.08 ± 0.06 |
| PEGDA(+)50 | 88.8 | 0 | + | 50 | 242.7 ± 0.38 | 0.09 ± 0.03 |
| PEGDA(-)50/dS | 68.8 | 20 | - | 50 | 674.5 ± 9.78 | 0.19 ± 0.02 |
| PS(-) | 0 | 0 | - | NA | 183.5 ± 1.1 | 0.03 ± 0.04 |
| PS(-)MP | 0 | 0 | - | NA | 1014.0 ± 27.0 | 0.14 ± 0.05 |

**Figure 1.**
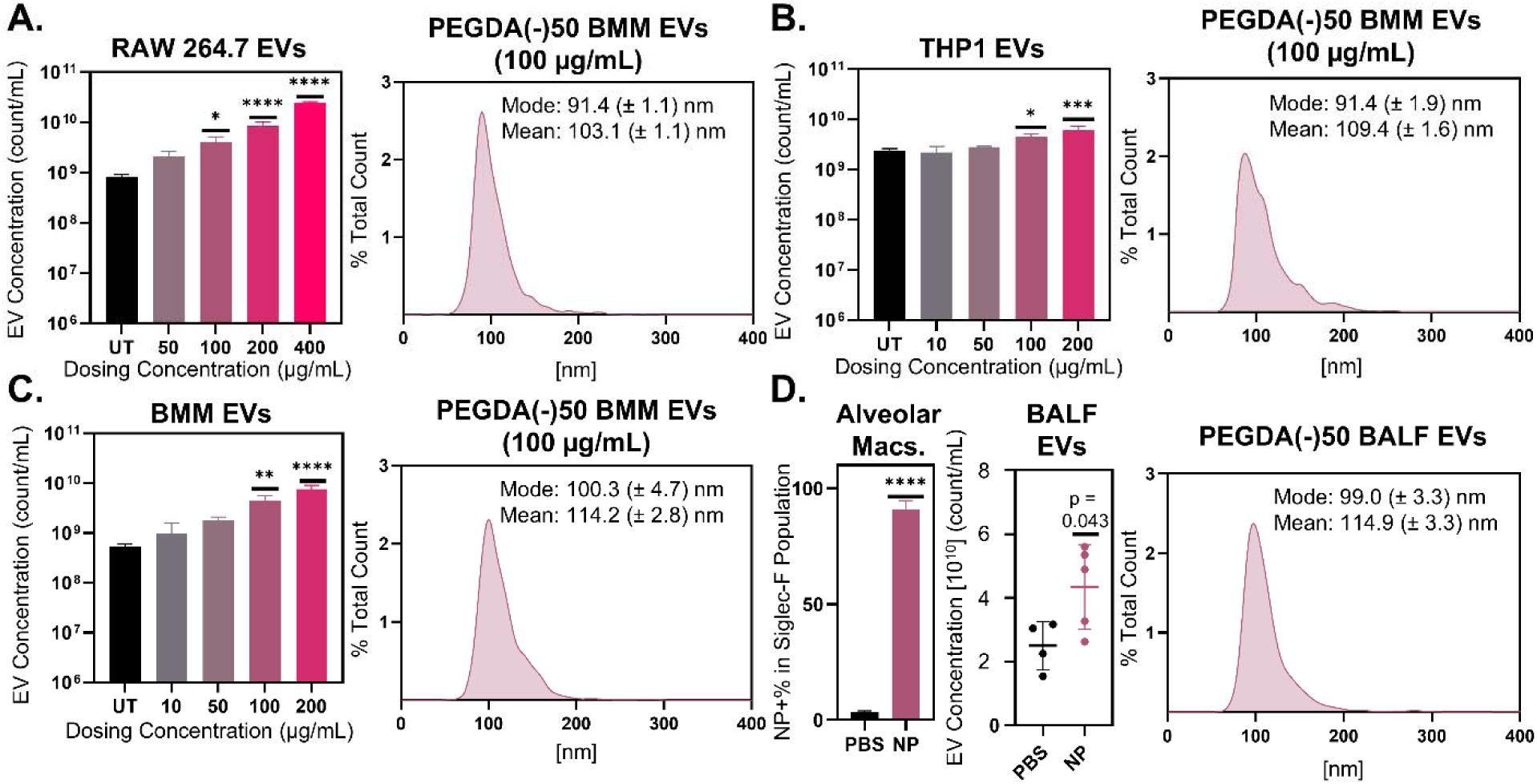
Particle internalization drives a dose dependent release of macrophage EVs. Nanoparticle tracking analysis (NTA) of ultracentrifuge isolated EVs as a variable of dosed PEGDA(-)50 for various cell types: **A)** RAW 264.7 macrophages immortalized and isolated from BALB/c mice, **B)** THP-1 immortalized human monocytes terminally differentiated into macrophages, **C)** *ex vivo* primary bone marrow derived macrophages (BMM) isolated from c57bl/6 mice (N = 3, SD error bars), *p<0.05, **p<0.01, ***p<0.001, ****p<0.0001 using ordinary one-way ANOVA with Turkey’s multiple comparisons test. **D)** Murine alveolar macrophages internalized PEGDA(-)50 orotracheal delivery (N = 4, SD error bars), ****p<0.0001 using unparied student’s T-test. **E)** NTA of ultracentrifuge isolated EVs from murine BALF (N = 4, SD error bars), p = 0.043 using unpaired student’s T-test. EVs isolated 72 hrs after particle dosing via differential centrifugation in all studies.

This effect was replicated in the human macrophage THP-1 cell line, in which THP-1 monocytes were terminally differentiated into macrophages prior to experimentation. As shown in **Figure 1B**, THP-1 macrophage *in vitro* cultures were dosed with 10 to 200 µg/mL of PEGDA(-)50, a lower concentration range relative to RAW 264.7 cells to account for their non-proliferative state. **Supplemental Figure S1.II** confirms PEGDA(-)50 internalization following 24 hrs in THP-1. Cultures that received 100 µg/mL exhibited approximately a 2-fold increase in ultracentrifugally isolated EV secretion collected 72 hrs after dosing compared to untreated cultures, while THP-1 macrophage cultures that received the highest dose (200 µg/mL) showed around a 2.5-fold increase in EV secretion (p = 0.0112 and p = 0.0001, respectively, via ordinary one-way ANOVA with Turkey’s multiple comparisons test). The size distribution for EVs from THP-1 cultures dosed with 100 µg/mL PEGDA(-)50 shows a similar size to EVs from RAW 264.7 dosed cultures and literature results, with a mode of 91.4 ± 1.9 nm and a mean of 109.4 ± 1.6 nm. The similarity of the RAW 264.7 and THP-1 macrophages points to a shared mechanism in engineered murine and human macrophages resulting in increased secretion of EVs following PEGDA(-)50 dosing.

We next performed similar studies using *ex vivo* macrophage cultures. In **Figure 1C**, primary-like bone marrow derived macrophages (BMMs) were isolated and differentiated per standard protocols^28^ and were dosed with 10 to 200 µg/mL of PEGDA(-)50. **Supplemental Figure S1.III** confirms PEGDA(-)50 internalization following 24 hrs in BMM and EVs were collected at 72 hrs. BMM cultures that received 100 µg/mL showed around an 8-fold increase in EVs compared to untreated (UT) cells, while the BMM cultures that received the highest concentration of 200 µg/mL showed a 14-fold increase in EV secretion (p = 0.0018 and p<0.0001, respectively, via ordinary one-way ANOVA with Turkey’s multiple comparisons test). Furthermore, the mode and mean of the 100 µg/mL dosed BMM cultures again mirrored those found in literature, despite being slightly larger than the cell line examples (100.3 ± 4.7 nm and 114.2 ± 2.8 nm, respectively).^29^ These results further demonstrate a direct relationship between PEGDA(-)50 dosing and EV secretion that is replicated in primary-like murine macrophages.

To confirm this effect was not an artifact of cell culture, we administered NPs to the lung space, where they should be mainly internalized by airway alveolar macrophages (AMs),^30^ and assessed changes to the airway EV profile. Using orotracheal delivery, we dosed healthy C57BL/6 mice with PBS (mock) or 100 µg of PEGDA(-)50 and collected bronchoalveolar lavage fluid (BALF) 72 hrs after dosing to measure changes in EV concentration. AM Siglec-F+ populations showed a significant increase in NP-derived Cy5 for mice dosed with PEGDA(-)50s (**Figure 1D**), confirming successful dosing of PEGDA(-)50 to AMs (representative gating for BALF-isolated AMs in **Supplemental Figure S2**). EV concentrations showed an increase in BALF EVs from mice that received PEGDA(-)50s as compared to the mock (p = 0.043 via unpaired Student’s T-test), with an EV size mean and mode of 99.0 ± 3.3 nm and 114.9 ± 3.3 nm that is consistent with prior *in vitro* results. While BALF EVs represent a heterogeneous population from multiple airway cell types, the strong NP uptake by Siglec-F+ AMs makes macrophages the most plausible source of the increased EV abundance. Thus, although macrophage-specific EV isolation from the BALF was not performed, these data support the conclusion that PEGDA(-)50-dependent EV response also occurs *in vivo* following dosing.

### Particle lysosomal accumulation is the main driver of EV secretion following phagocytosis

Having demonstrated that PEGDA(-)50 NPs drove macrophage EV secretion in various *in vitro* cultures and *in vivo*, we sought to evaluate the driving particle characteristics responsible for EV secretion (see complete particle chemistries and characterization in **Table 1**). To screen for the role of surface charge, particle chemistry, and particle size, EV concentrations were first compared between BMMs dosed with equivalent mass (100 µg/mL) of anionic PEGDA(-)50 and cationic PEGDA(+)50, as well as polystyrene (PS) NPs [PS(-)] ∼180 nm in diameter and PS microparticles [PS(-)MP] ∼1 µm in diameter. **Figure 2A** shows that particle chemistry or charge does not drive differences in particle amplified EV secretion. Collected 72 hrs after dosing, BMM cultures secreted more EVs following dosing with PEGDA(-)50, PEGDA(+)50, and PS(-)MP as compared to UT (p = 0.0006, p = 0.0002, and p = 0.0316, respectively, via ordinary one-way ANOVA with Turkey’s multiple comparisons test). The EVs amplified by distinct particles had similar size distributions: modes of 100.3 ± 4.7 nm (**Figure 1C**), 92.6 ± 1.8 nm (**Figure 2B.i**), and 100.3 ± 1.7 nm (**Figure 2B.ii**) for PEGDA(-)50, PEGDA(+)50, and PS(-)MP, respectively, with representative TEM images of ultracentrifugally isolated PEGDA(-)50 amplified BMM EVs shown in **Figure 2C**. Interestingly, PS(-), the smallest particle size tested at ∼180 nm, did not increase secreted EVs when dosed at equivalent mass to the larger particles, nor did treatment with lipopolysaccharide (LPS), a macrophage activation control. Together, these findings show that particle charge and chemistry do not regulate the EV increase, but rather a critical particle size is crucial in driving this effect.

**Figure 2.**
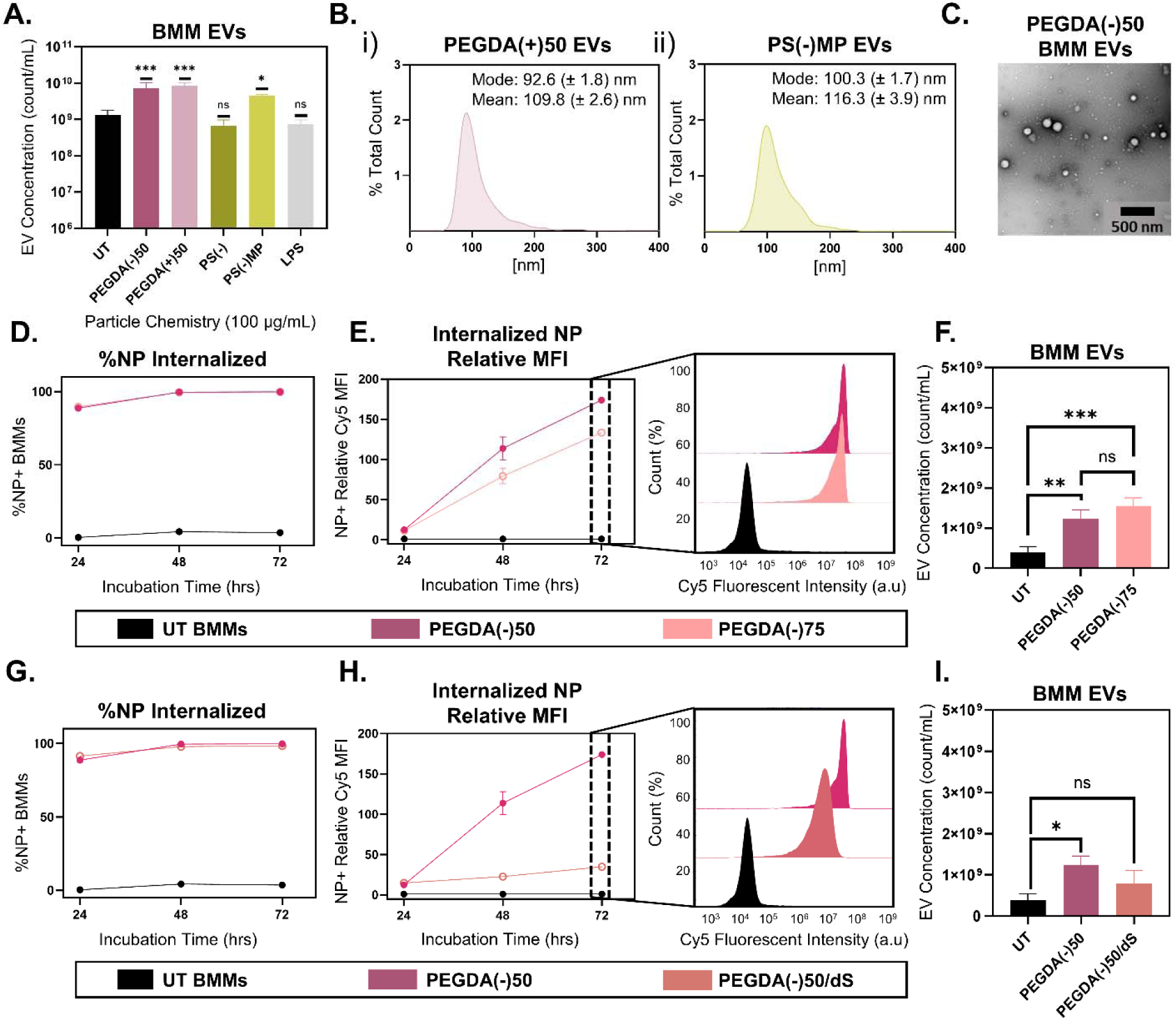
Particle physiochemical properties alter EV secretion in BMMs. **A)** NTA analysis of ultracentrifuge isolated BMM EVs 72 hrs following dosing with 100 µg/mL of various particle chemistries (PEGDA(-)50, PEGDA(+)50, PS(-), and PS(-)MP) and LPS, (N = 3, SD error bars), *p<0.05, ***p<0.001, using ordinary one-way ANOVA with Turkey’s multiple comparison test. **B)** NTA size distribution for PEGDA(+)50 and PS(-)MP amplified BMM EVs (i. and ii. respectively). **C)** TEM scans of isolated PEGDA(-)50 amplified BMM EVs (500 nm scale bar). **D)** Percent NP+ population of BMMs following 24, 48, and 72 hrs after dosing with PEGDA(-)50 and PEGDA(-)75 (N = 3). **E)** Relative MFI for NP+ BMM populations 24, 48, and 72 hrs following dosing with PEGDA(-)50 and PEGDA(-)75, alongside 72 hrs BMM fluorescent intensity distribution (N = 3). **F)** NTA analysis of ultracentrifuge isolated BMM EVs 72 hrs following dosing with 100 µg/mL of PEGDA(-)50 and PEGDA(-)75, **<0.01, ***<0.001, using ordinary one-way ANOVA with Turkey’s multiple comparison test (N = 3). **G)** Percent NP+ population of BMMs following 24, 48, and 72 hrs after dosing with PEGDA(-)50 and PEGDA(-)50/dS (both line present, PEGDA(-)50/dS line behind PEGDA(-)50 line, N = 3). **H)** Relative MFI for NP+ BMM populations 24, 48, and 72 hrs following dosing with PEGDA(-)50 and PEGDA(-)50/dS, alongside 72 hrs BMM fluorescent intensity distribution (N = 3). **I)** NTA analysis of ultracentrifuge isolated BMM EVs 72 hrs following dosing with 100 µg/mL of PEGDA(-)50 and PEGDA(-)50/dS, **<0.01, ***<0.001, using ordinary one-way ANOVA with Turkey’s multiple comparison test (N = 3).

To distinguish whether EV amplification depends on the number of particles internalized or the total mass delivered per cell, we compared PEGDA(-)50 to a denser formulation at 75 wt% [PEGDA(-)75] dosed at equal mass to BMMs. This design allowed us to vary particle number while keeping delivered mass constant (i.e., the PEGDA(-)50 will have a greater number than the PEGDA(-)75 at an equivalent mass dose, **Supplemental Figure S3**). Across both formulations, more than 80% of dosed BMMs were positive for particle internalization within 24 hrs, and this uptake level remained stable across the 72 hrs incubation period used for EV isolation (**Figure 2D**). However, Cy5 median fluorescent intensity (MFI) increased over time to different degrees; PEGDA(-)75 showed lower accumulation at 72 hrs than the PEGDA(-)50 (**Figure 2E**), consistent with fewer particles per cell at equal mass. Interestingly, despite these differences, **Figure 2F** shows that both the PEGDA(-)50 and PEGDA(-)75 induced a similar EV amplification response (p = 0.004 and p = 0.0007, respectively, via ordinary one-way ANOVA with Turkey’s multiple comparison test) compared to UT. These results indicate that EV secretion scales with total particulate mass delivered per cell, rather than the number of individual phagocytic events.

To test whether EV amplification depends on the intracellular residence time of internalized particles, we compared the standard PEGDA(-)50 formulation to a fast-degrading PEGDA(-)50/dS chemistry designed to break down within lysosomes.^27^ In both formulations, over 80% of dosed BMMs were positive for particle internalization through 72 hrs (**Figure 2G**). In contrast to results from **Figure 2E**, the fast-degrading PEGDA(-)50/dS chemistry showed minimal increase in MFI over the same period (**Figure 2H**), as highlighted by the fluorescent intensity histograms compared to the PEGDA(-)50 chemistry at 72 hrs. Correspondingly, **Figure 2I** shows that, while the PEGDA(-)50 amplified the EV secretion of BMMs, the fast-degrading PEGDA(-)50/dS failed to drive EV amplification of BMMs (p = 0.1877 via ordinary one-way ANOVA with Turkey’s multiple comparison test). Together, these findings indicate that prolonged lysosomal accumulation and intracellular processing are required to trigger the EV amplification response.

### Electron microscopy captures the formation of MVB following particle dosing

To determine whether increased EV secretion arose from surface budding (i.e., microvesicle formation) or from intracellular pathways, we first examined cell morphology by scanning electron microscopy (SEM) 24 hrs after particle dosing (100 µg/mL). **Figure 3** shows representative images of cells dosed with PEGDA(-)50, PEGDA(+)50, or PS(-)MP; red arrows indicate potential cell membrane blebbing, which would lead to the formation of microvesicles. **Supplemental Figure S4** depicts a second set of images with regions of interest. Across all conditions, we observed no increase in membrane protrusions or blebs that would indicate enhanced microvesicle shedding, as the frequency of membrane blebbing was constant across conditions. Notably, the PS(-)MP images captured active phagocytosis of the PS particles (**Figure 3D.ii**) as indicated by the yellow arrows. The absence of increased surface budding across all particle types suggests that the EV amplification observed in BMMs does not arise from microvesicle production and instead points toward intracellular mechanisms of EV biogenesis.

**Figure 3.**
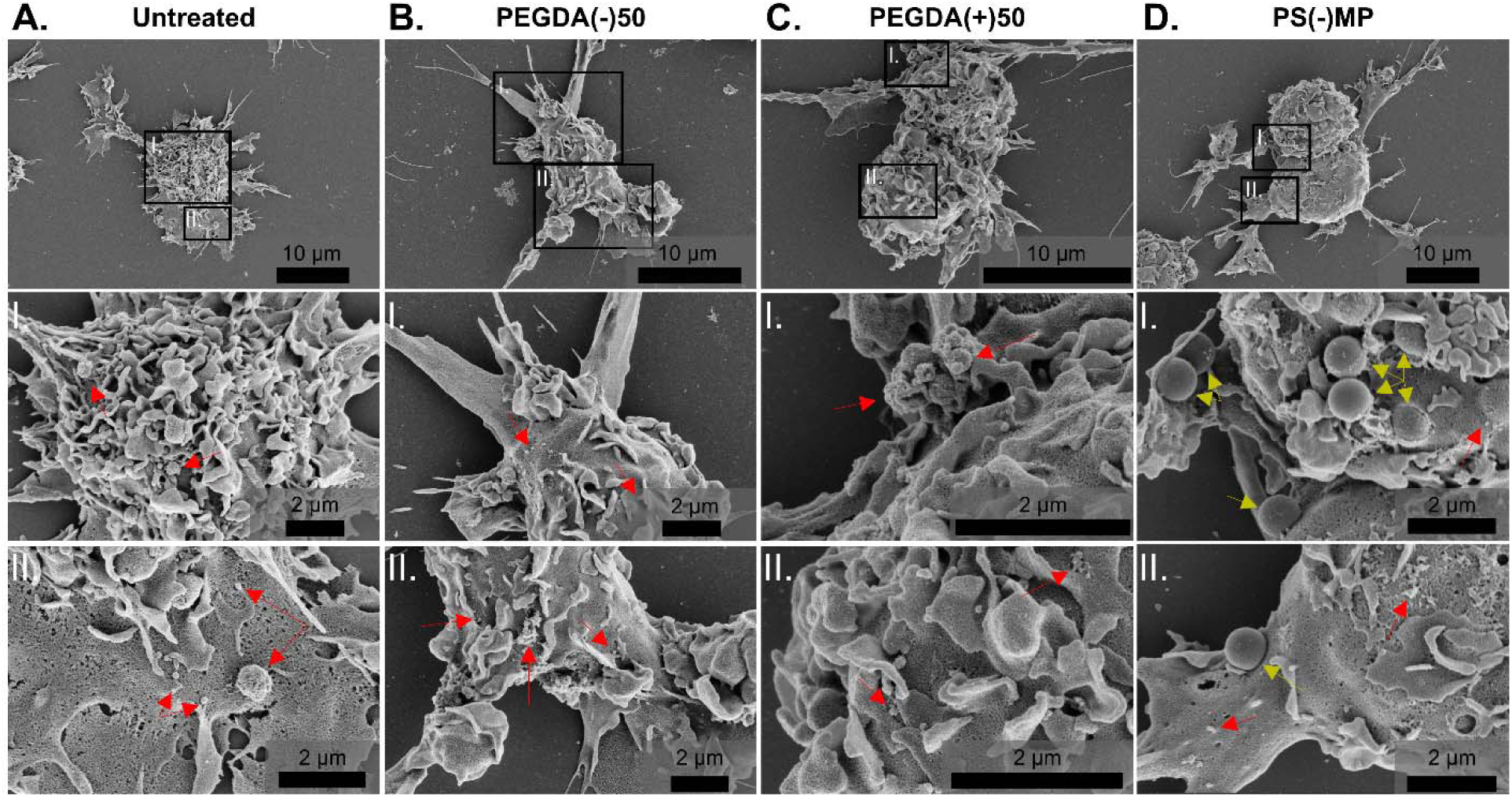
SEM imaging of particle-dosed BMM does not show an increase in surface budding. Selected BMM conditions taken 24 hrs after dosing were chosen for SEM imaging. **A)** Untreated, **B)** PEGDA(-)50, **C)** PEGDA(+)50 and **D)** PS(-)MP dosed-BMMs (scale bar 10 µm) with corresponding magnified sections (**I.** and **II.**, scale bar 2 µm). Red arrows indicate potential instances of membrane budding, yellow arrows indicate PS(-)MP on the cell surface. Additional images in **Supplemental Figure S4**.

To gain better insight as to these internal cellular structures, we performed TEM imaging of untreated, PEGDA(-)50, PEGDA(+)50, or PS(-)MP under the same conditions. **Figure 4** shows particles internalized into the lysosomal compartment (dark stain) of the cells, indicated by the white arrows labeled as NP. **Supplemental Figure S5** depicts a second set of images with regions of interest. Particle-treated cells showed a clear increase in the vesicles per cell, **Supplemental Figure S6**. Furthermore, of these vesicles, more were MVBs, the intracellular structures that generate small EVs, relative to untreated cultures.^31^ These MVBs were frequently observed near internalized particles and co-localized near lysosome. This spatial association suggests that particle processing in the lysosome promotes MVB biogenesis, linking phagocytosis to enhanced EV release.

**Figure 4.**
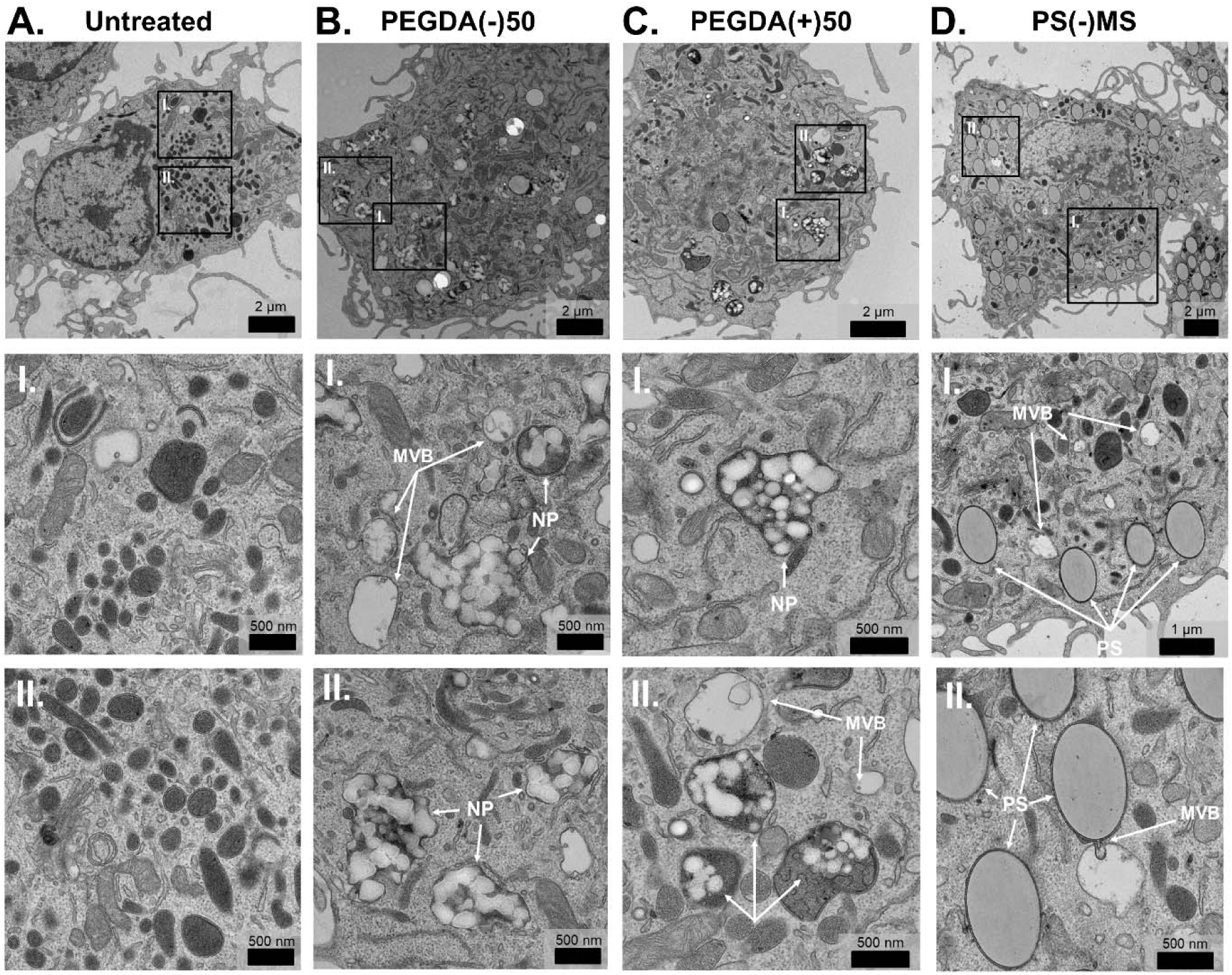
TEM imaging of particle dosed *ex vivo* BMM shows MVB formation and lysosomal co-localization after 24 hrs. Selected BMM conditions 24 hrs after dosing were chosen for TEM imaging. **A)** Untreated, **B)** PEGDA(-)50, **C)** PEGDA(+)50 and **D)** PS(-)MP dosed-BMMs (scale bar 2 µm) with corresponding magnified sections (**I.** and **II.**, scale bar 500 nm). MVB and NPs / PS are labelled in insets. Additional images in **Supplemental Figure S5**.

### Gene enrichment analysis of proteomic data indicates increased vesicle formation and metabolic activity

For further insight into the intracellular mechanism following NP internalization, we performed proteomic analysis of the cellular and secreted proteome of BMMs dosed with 100 µg/mL of PEGDA(-)50. Whole-cell proteomics were performed at 24 hrs to characterize early intracellular responses following NP internalization, while secreted proteomics were collected at 48 hrs to ensure adequate accumulation of released EVs in the supernatant. Secreted proteomic analysis was completed on all secreted signals and not on isolated BMM EVs. **Figure 5A** shows the quantity of proteins that were statistically enriched (524 proteins) or decreased (527 proteins) in the cellular proteome following PEGDA(-)50 dosing (Student’s T test, False Discovery Rate, FDR = 0.1). At the same time, **Figure 5B** shows the quantity of proteins that were statistically enriched (136 proteins) or decreased (77 proteins) in the secreted proteome following PEGDA(-)50 dosing (Student’s T test). **Supplemental Figure S7** shows the respective volcano plots. Hierarchical heat maps of the identified differentially expressed proteins in the whole-cell and secreted proteomics datasets (FDR = 0.1) are shown in **Figure 5C** and **5D**, respectively. We note that the magnitude of differential expression is small, with the average z-score for significantly differentially expressed proteins identified in both whole-cell and secreted datasets being less than 1. This minimal change in protein composition is expected based on previously published work,^28^ which showed that this negatively charged PEGDA(-)50 chemistry had minimal effect on the phenotype of BMMs. **Supplemental Figure S8** shows a hierarchical clustering heat map for UT, PEGDA(-)50, and pro-inflammatory control LPS to confirm the lack of macrophage phenotype activation in PEGDA(-)50-treated BMMs.

**Figure 5.**
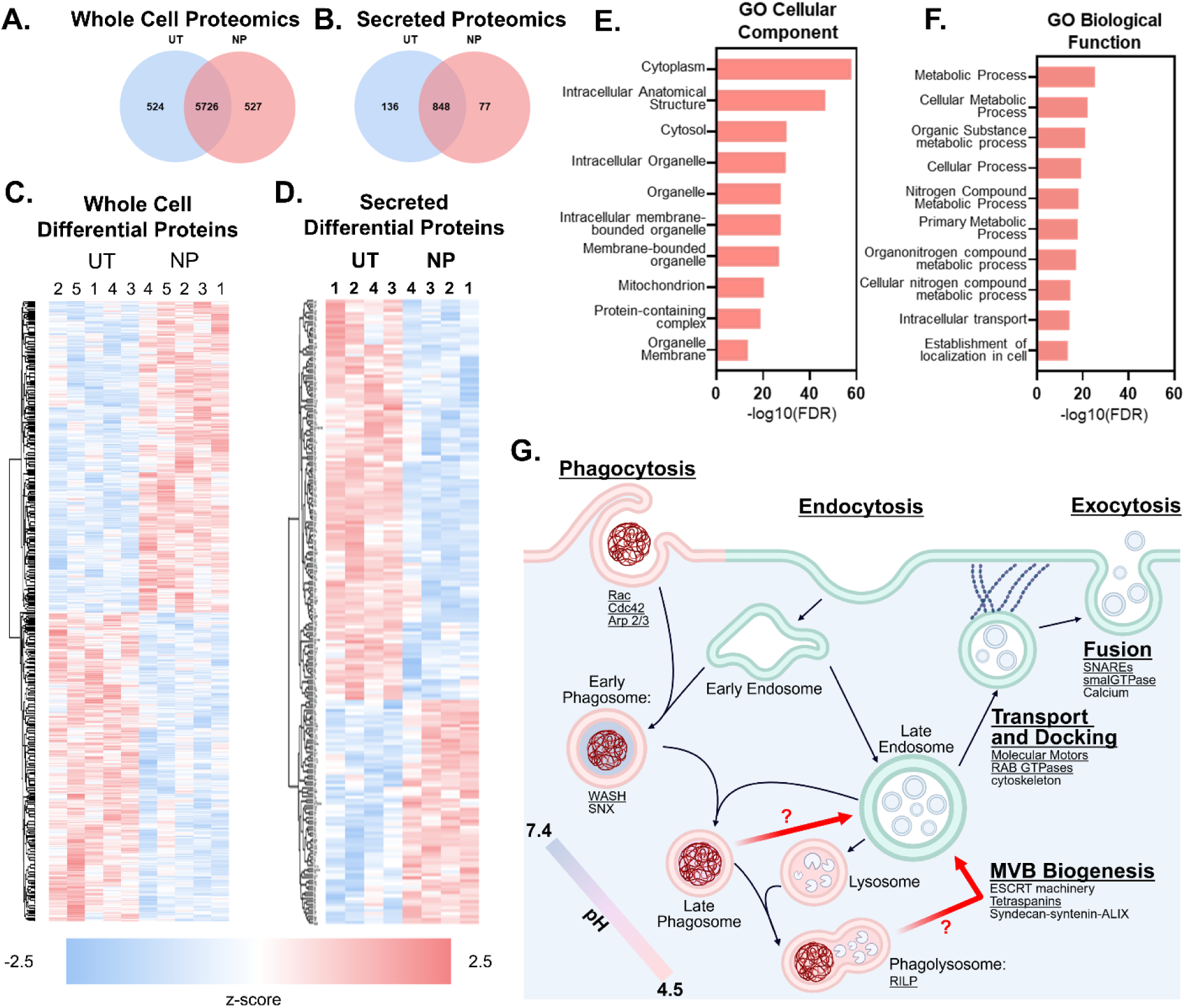
Proteomic analysis of PEGDA(-)50 (NP) dosed BMM after 24 hours shows enrichment for metabolic activity and membrane-bounded organelles. Significantly differentiated proteins for proteomic analysis of **A)** whole cell proteome (N = 5) and **B)** secreted proteome (N = 4) show identification of 1051 and 213 hits, respectively (Student’s T-test, FDR = 0.1). Hierarchical clustering of significantly differentiated proteins for **C)** whole cell proteomics and **D)** secreted proteomics. Performed gene enrichment analysis of clusters with increased presence in NP dosed conditions. **E)** Gene enrichment analysis using GO cellular component shows a significant increase for membrane-bounded organelles. **F)** Enrichment analysis performed with GO biological function as the database shows an increase in metabolic process following dosing with NP. **G)** Schematic of hypothesized phagocytosis driven EV biogenesis pathway; underlined proteins are upregulated in PEGDA(-)50 dosed BMMs.

Following identification of upregulated species throughout the datasets, we performed gene enrichment analysis to identify possible pathways with increased activity following dosing with PEGDA(-)50. Because our focus was on vesicle trafficking and structural pathways, we examined enriched terms within the Gene Ontology (GO) Cellular Component database (**Figure 5E**).^32, 33^ Among the top categories, several were associated with membrane-bound compartments (*i.e.*, intracellular anatomical structure, intracellular organelle, etc.), and proteins increased in the secreted dataset showed an enrichment for extracellular exosomes (FDR = 0.0358). Together with prior data shown in Figures 1-4, these proteomic signatures support a model in which particle phagocytosis promotes small EVs through the enhanced MVB formation.

We also evaluated broader functional changes using GO Biological Process analysis. Several enriched pathways reflected shifts in cellular metabolism following PEGDA(-)50 dosing (**Figure 5F**), with five of the top ten terms related to metabolic processes.^32, 33^ Although the associated FDR values are modest, these subtle metabolic changes are consistent with prior observations for this particle chemistry reported to drive increased lysosomal activity and cell survival.^28^ **Supplemental Figure S9** shows, through a CellTiter Glo 2.0 assay, that BMMs dosed with PEGDA(-)50 NPs have increased metabolic activity, corroborating the gene enrichment result. Such changes could contribute to, or arise alongside, the increased MVB activity underlying small EV biogenesis.

Based on this analysis, in line with relevant literature,^14^ we have hypothesized a pathway for phagocytosis-driven EV biogenesis (**Figure 5G**). Canonical phagocytosis structures are delineated by the red lipid membrane, while canonical small EV biogenesis structures are delineated by the green lipid membrane. Critical proteins and protein families for these pathways are noted in the schematic, with proteins or protein families that were found to have increased presence in PEGDA(-)50-dosed BMM in our proteomic studies are underlined for emphasis. The pH of phagocytosis-associated vesicles is indicated by the color inside the vesicle structure based on established literature.^34^ Critical to this work, we hypothesize the existence of a connecting pathway between the phagocytosis process and the small EV biogenesis process, indicated by red arrows. Based on our data, we hypothesize two potential connection points between these processes: 1) diversion of late phagosomes to MVBs in response to material accumulation in the phagolysosome, or 2) transfer of degraded material from the phagolysosome to an MVB.

### Source macrophage phenotype shapes cargo and transfer properties of NP-amplified EVs

EV cargo is strongly shaped by the activation state of the EV-producing cell;^5, 6, 35^ we therefore examined how producing cell (i.e., “source” BMM) phenotype influences the composition and function of particle-amplified EVs. BMMs were polarized alongside PEGDA(-)50 dosing 24 hrs after initial seeding using historically defined macrophage activation states representing canonical extremes along a broader phenotypic spectrum: “M0” inactivated, “M1” classically-activated by 20 ng/mL IFN-γ, and “M2” alternatively-activated by 20 ng/mL IL-13. Phenotype of these polarized cells 72 hrs following stimuli was confirmed in **Supplemental Figure S10** and **Supplemental Figure S11** for pro-inflammatory and anti-inflammatory, respectively. To rapidly characterize the resultant EV composition, we used a commercially available streptavidin-coated magnetic bead assay using biotinylated capture antibodies to identify isolated BMM EVs of varied phenotypes and characterize their membrane protein expression (**Figure 6A**). **Figure 6B** shows the production timeline for characterized BMM EVs, where incubation time (T_I_) is the time between initial PEGDA(-)50 dosing and BMM-EV isolation. **Figure 6C** depicts the three components characterized in this study: source BMMs, their secreted BMM-EVs, and naïve BMM EV-recipients. For convenience, the BMM phenotype (M0, M1, or M2) is indicated on the label with the treatment type (NP or EV) as a subscript. For EVs, the dosing condition of the source BMM (M0, M0-NP, M1-NP, M2-NP) is represented as a superscript.

**Figure 6.**
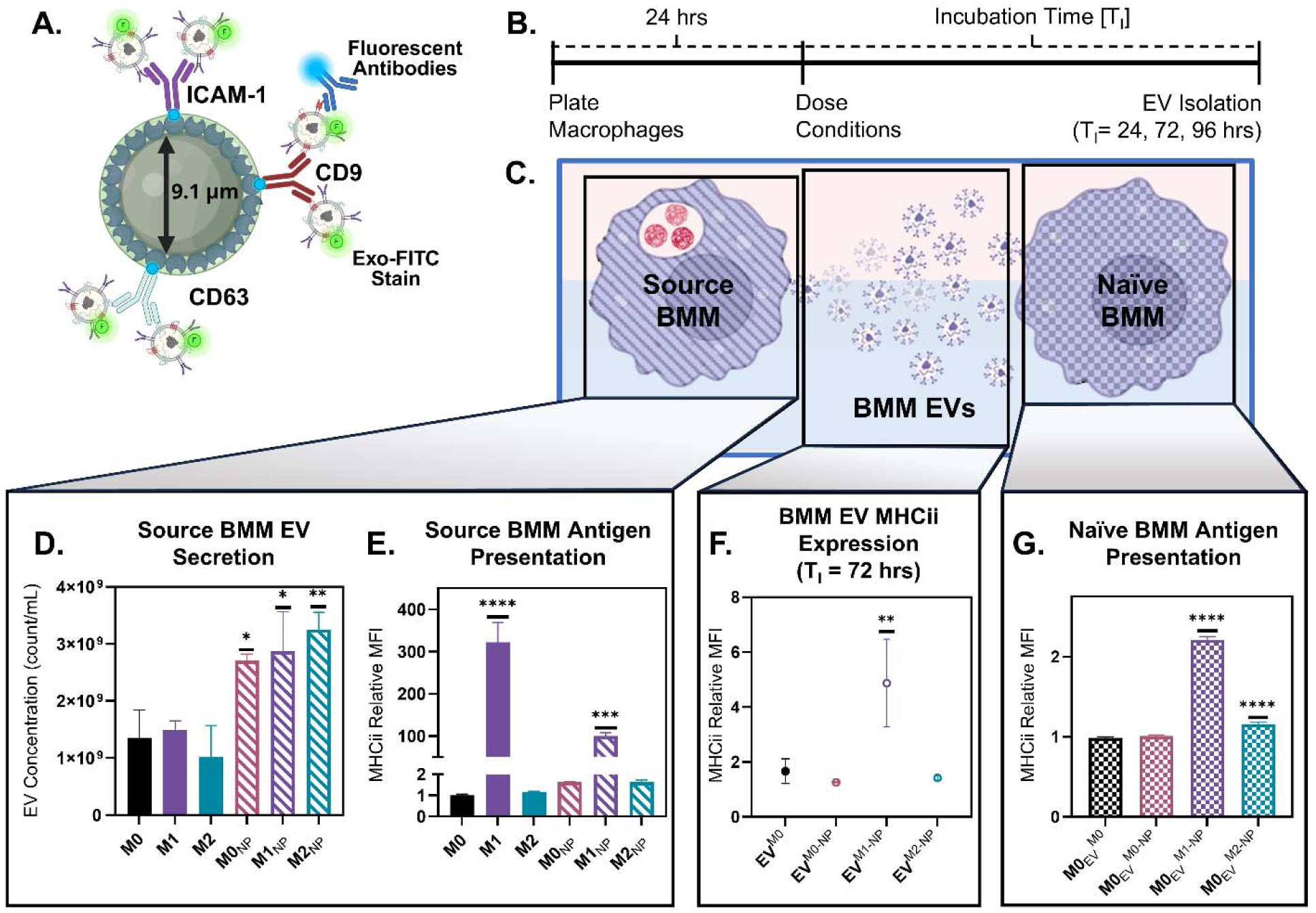
PEGDA(-)50 (NP)-amplified EVs transfer antigen-presenting signaling depending on source cell phenotype. **A.**) Schematic of modular magnetic bead assay for flow cytometry characterization of isolated EV stocks from BMM cells using anti-CD63, CD9, and ICAM-1 antibodies as capture antibodies. **B)** EV isolation timeline for EV stock characterization; incubation time (T_I_) indicating the time between initial dosing of cells with NP and EV isolation. **C)** Schematic depicting macrophage to macrophage transfer of cell signal via EVs following dosing with NP. For the following dosing conditions, naming convention are as following: [BMM Phenotype]_[NP or EV dosing]_^[Source BMM Phenotype]^. **D)** NTA analysis of ultracentrifuge isolated BMM EVs for phenotypically active macrophages (M0: inactivated macrophages, M1: pro-inflammatory, 20 ng/mL IFN-γ, M2: anti-inflammatory, 20 ng/mL IL-13) and phenotypically active macrophages dosed with PEGDA(-)50, *<0.05, **<0.01, using ordinary one-way ANOVA with Turkey’s multiple comparison test (N = 3). **E)** Flow cytometry analysis of MHCii expression as a marker for antigen presentation in phenotypically active macrophages and phenotypically active macrophages dosed with PEGDA(-)50 after T_I_ = 72 hrs, ***<0.001, ****<0.0001, using ordinary one-way ANOVA with Turkey’s multiple comparison test (N = 3), relative to M0. **F)** Anti-CD63 capture bead flow cytometry analysis of MHCii presence in isolated BMM EVs for M0 macrophages and M0, M1, and M2 macrophage dosed with PEGDA(-)50, **<0.01, using ordinary one-way ANOVA with Turkey’s multiple comparison test (N = 3). **G)** Flow cytometry analysis of MHCii expression in naive macrophages dosed with EVs from M0 macrophages and M0, M1, and M2 macrophage dosed with PEGDA(-)50, ****<0.0001, using ordinary one-way ANOVA with Turkey’s multiple comparison test (N = 3), relative to M0_EV_^M0^.

We first examined whether the BMM phenotype influences EV secretion in the absence of particles. Untreated M0, M1, and M2 macrophages released comparably consistent EV levels, indicating that activation does not substantially enhance EV output. Upon PEGDA(-)50 dosing, all phenotypes showed a robust EV increase (**Figure 6D**; M0_NP_(p-value) = 0.0254, M1_NP_(p-value) = 0.0118, and M2_NP_(p-value) = 0.0021 via ordinary one-way ANOVA with Turkey’s multiple comparisons), demonstrating that phenotype does not dictate EV secretion. As a representative marker, we assessed MHC II expression in polarized cells, as it is a canonical antigen-presentation marker known to load onto immune cell-derived EVs.^14, 36^ All conditions were normalized to the M0 condition. As expected, M1(IFN-γ) BMMs expressed significantly higher MHC II than M0 or M2 cells, and this hierarchy was maintained following PEGDA(-)50 treatment (**Figure 6E**; M1(p-value) < 0.0001, and M1_NP_(p-value) = 0.0005 compared to M0 MHC II expression, via ordinary one-way ANOVA with Turkey’s multiple comparisons). These results further indicate that PEGDA(-)50 amplified EV secretion without altering the underlying polarization phenotype of the source macrophages.

Having established that PEGDA(-)50 does not alter macrophage polarization state, we next asked whether the preserved source-cell phenotype is imprinted onto the released EVs and whether these phenotype-specific signatures can transfer to naïve macrophages. Using the magnetic beads depicted in **Figure 6A** with anti-CD63 capture antibodies, we characterized MHC II expression in various isolated EV stocks: EVs from M0 BMMs (EV^M0^), EVs from M0 BMMs dosed with PEGDA(-)50 (EV^M0-NP^), EVs from M1 BMMs dosed with PEGDA(-)50 (EV^M1-NP^), and EVs from M2 BMMs dosed with PEGDA(-)50 (EV^M2-NP^). **Figure 6F** shows that the EV^M1-NP^ had an increased expression of MHC II relative to the EV^M0^ (p = 0.0064, compared to EV^M0^, via ordinary one-way ANOVA with Turkey’s multiple comparison test), mirroring the expression profile shown in **Figure 6E**. Finally, ultracentrifugally isolated BMM-EVs from various phenotypes were dosed (1*10^9^ count/mL) onto naïve M0 BMMs to determine the potential for EV-mediated transfer of host cell expression. **Figure 6G** shows that both M0 BMMs dosed with EVs from M1_NP_ (M0_EV_^M1-NP^) and M0 BMMs dosed with EVs from M2_NP_ (M0_EV_^M2-NP^) exhibited increased MHC II expression (p < 0.0001 and p = 0.0002, respectively, compared to M0_EV_^M0^, via ordinary one-way ANOVA with Turkey’s multiple comparison test). Interestingly, the slight but significant increase in MHC II expression for M0_EV_^M2-NP^ does not mirror the MHC II expression characterized in **Figure 6F** (EV^M2-NP^). This could point to the importance of other encapsulated biological signals, such as RNA, that this flow cytometry-based characterization would not capture.^37^ In combination, these findings show that by controlling the host source macrophage phenotype, there is the potential to generate macrophage-EVs with unique phenotypes at amplified quantities, which can be transferred to other naïve macrophages.

### Particle material is loaded in phagocytosis-driven EVs and transferred to naïve macrophage cells

Since particles drive the amplification of macrophage-EV secretion, we next asked whether particle material itself is transferred into these EVs. To confirm the transfer of particle material, we tracked the Cy5 fluorophore covalently attached to the PEGDA(-)50 and quantified its presence in varied EV samples. We first used anti-CD63 magnetic beads to measure Cy5 MFI for RAW 264.7 macrophages following dosing with 100 µg/mL of PEGDA(-)50 and a T_I_ of 24, 72, or 96 hrs. **Figure 7A** shows that the highest and most consistent Cy5 signal occurred at 24 hrs (p = 0.0229, via unpaired t-test with Welch’s correction). Although the Cy5 signal was still detectable at 72 hrs, the degree of transfer varied significantly between replicates (**Supplemental Figure S12**), suggesting temporal heterogeneity in NP internalization, digestion, and EV loading. Cy5 was covalently incorporated into the PEGDA(-)50 network during polymerization; thus, its appearance on isolated EVs indicates that at least a portion of the phagocytosed particles must have undergone intracellular degradation to release Cy5-containing fragments that can be packaged into EVs, confirming that particle-derived material was transferred into secreted EVs.

**Figure 7.**
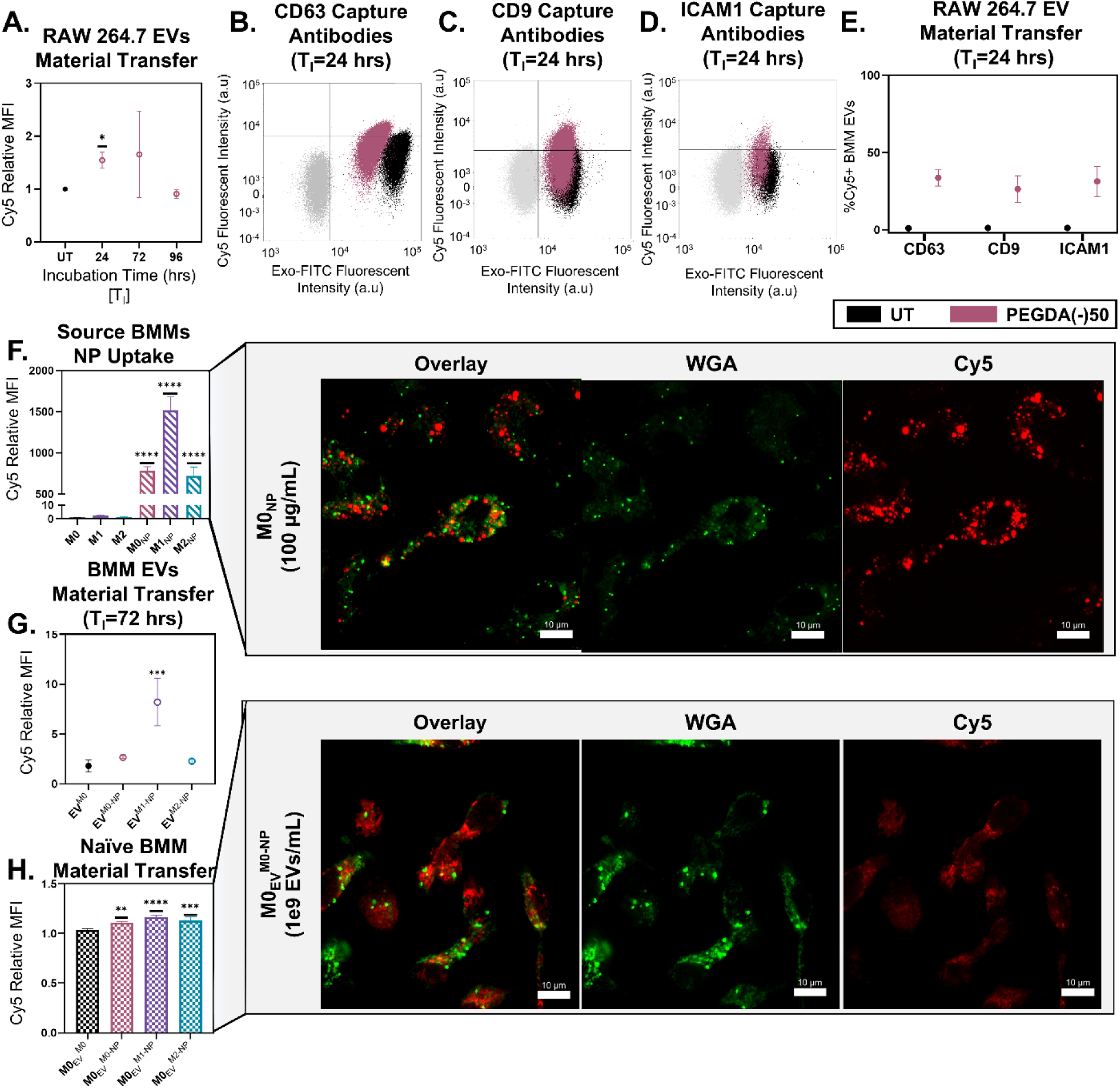
PEGDA(-)50 (NP)-amplified EVs transfer NP material from source to receiving macrophages. **A)** Anti-CD63 capture bead flow cytometry analysis of relative Cy5 MFI in PEGDA(-)50 amplified EVs from RAW264.7 cells as a marker for NP material transfer for T_I_ = 24, 48, and 72 hrs, *p<0.05, via unpaired t-test with Welch’s correction (N = 3). Dot plot of capture antibody flow cytometry analysis of EVs from RAW264.7 cells at 24 hrs for **B)** anti-CD63, **C)** anti-CD9, and **D)** anti-ICAM-1, showing bead control population (gray), UT EV population (black), and PEGDA(-)50 amplified EVs (pink). **E)** %Cy5^+^ population for UT (black) and PEGDA(-)50 amplified EVs from RAW264.7 cells (pink) using anti-CD63, CD9, and ICAM-1 capture antibodies, via ordinary two-way ANOVA with Bonferri multiple comparisons test (N = 3). **F)** Flow cytometry analysis of relative Cy5 MFI in phenotypically active macrophages and phenotypically active macrophages dosed with PEGDA(-)50 after T_I_ = 72 hrs, ****<0.0001, using ordinary one-way ANOVA with Turkey’s multiple comparison test (N = 3), and **i)** associated confocal imaging using (scale bar 10 µm). **G)** Anti-CD63 capture bead flow cytometry analysis of MHCii presence in isolated BMM EVs for M0 macrophages and M0, M1, and M2 macrophage dosed with PEGDA(-)50, ***<0.001, using ordinary one-way ANOVA with Turkey’s multiple comparison test (N = 3). **H)** Flow cytometry analysis of relative Cy5 MFI in inactivated BMMs dosed with EVs from M0 macrophages and M0, M1, and M2 macrophage dosed with PEGDA(-)50, **p<0.01, ***<0.001, ****<0.0001, using ordinary one-way ANOVA with Turkey’s multiple comparison test (N = 3), and **i)** associated confocal imaging using (scale bar 10 µm).

To determine whether NP loading occurred across distinct subpopulations, we captured EVs using antibodies against CD63, CD9, and ICAM-1. CD63 and CD9 have been shown to transfer to the membranes of small EVs during the biogenesis process, while ICAM-1 is a targeting surface protein often found in EVs.^17^ **Figure 7B-D** shows an example flow cytometry dot plot of stained magnetic beads (gray), UT BMM EVs (black), and PEGDA(-)50 dosed BMM-EVs (red) at T_I_=24 hrs using anti-CD63, CD9, and ICAM-1 capture beads, respectively. Exo-FITC gates (indicating the presence of EVs on the beads) were set relative to the stained magnetic bead population, while the Cy5 gates were set relative to the UT EV population. **Figure 7E** shows the replicate results for each EV subpopulation, showing a consistent increase in Cy5+ populations, CD63(mean) = 26.83%, CD9(mean) = 31.31%, and ICAM1(mean) = 33.65% (p = 0.0025, p = 0.0005, and p = 0.002, respectively, via ordinary 2way ANOVA with Bonferroni’s multiple comparisons test). These results confirm that particle material transfers into diverse EV subpopulations, potentially with distinct EV protein expression.

Following this, we wanted to confirm that this particle material transfer occurs independently of host cell phenotype. We dosed 100 µg/mL of PEGDA(-)50 to BMMs of varied polarization, showing that PEGDA(-)50 internalization persists for 72 hrs (**Figure 7F**), normalized to the baseline Cy5 expression in the M0 condition. Consistent with the literature, M1_NP_ BMMs exhibited a significant increase in NP uptake compared to the M0_NP_.^38^ Representative confocal images of M0_NP_ show punctate Cy5 fluorescence (**Figure 7F inset**) that is co-localized mainly with WGA-stained membrane-bound compartments, indicating their presence within intracellular phagolysosome structures.^39^

Using CD63-beads to capture EVs, we observed a significant increase in Cy5 fluorescence for EVs collected from PEGDA(-)50-treated M1 macrophages (**Figure 7G**; EV^M1-NP^(p-value) = 0.0010, via ordinary one-way ANOVA with Turkey’s multiple comparisons test) normalized to the EV^M0^ condition. These data indicate that particle-material loading into EVs occurs even in phenotypically distinct BMM states. Interestingly, **Figure 7H** shows effective transfer of NP Cy5 material for all PEGDA(-)50 amplified EVs. Naïve M0 BMMs were dosed with EVs collected from polarized, PEGDA(-)50-treated BMMs at 1*10^9^ EV counts/mL, and the Cy5 in the recipient naïve cell was measured. Despite overall only slight increases, each condition that received PEGDA(-)50 amplified EVs was detected at a significant amount compared to the control M0_EV_^M0^–M0 BMMs receiving EVs from UT M0 BMMs (M0_EV_^M0-NP^(p-value) = 0.0109, M0_EV_^M1-NP^(p-value) = 0.0003, and M0_EV_^M2-NP^(p-value) = 0.0021 via ordinary one-way ANOVA with Turkey’s multiple comparisons). Representative confocal images of M0_EV_^M0-NP^ show diffuse Cy5 fluorescence (**Figure 7H inset**) that is present throughout the cytosol. This suggests a distinct internalization method for the EVs compared to the PEGDA(-)50 particle itself.

## DISCUSSION

Despite the ubiquity of phagocytosis in macrophage biology, the idea that particulate phagocytosis directly regulates small EV biogenesis has not previously been demonstrated. Prior work has generally attributed EV increases to broad cellular stress responses, leaving open the question of whether particulate burden itself can regulate EV formation.^40, 41^ In this work, we demonstrate that phagocytic uptake and intracellular trafficking of particulate materials act as primary drivers of small EV secretion, directly bridging the canonical understanding of phagocytosis and EV biogenesis pathways for the first time.

Phagocytosis stands as one of the crucial homeostatic processes and is a subclass of endocytic pathways. Engagement of cell-surface receptors with target particles drives actin-dependent membrane remodeling and formation of phagosomes.^42, 43^ As phagosomes mature, they acquire endosomal and lysosomal components, undergo progressive acidification, and ultimately fuse with lysosomes to form phagolysosomes that degrade internalized material.^13, 42^ The formation of the phagolysosome is often seen as the endpoint of this process, where material is degraded, and byproducts are released via aqueous efflux.^34^ The phagosome maturation process parallels the endocytosis pathway involved in small EV biogenesis.^44^ During small EV biogenesis, early endosomes mature into a subcategory of late endosomes called MVBs, characterized as vacuoles with smaller intraluminal vesicles (ILV) inside. Following this maturation, MVBs transport their cargo to the cell membrane, docking it via a plethora of molecular motors,^45^ RAB GTPases,^46^ and cytoskeletal proteins.^47^ SNARE^48^ and small GTPases^49^ then facilitate the fusion of these vacuoles and the cell membrane, resulting in the release of the ILVs into the extracellular space, *i.e.,* exocytosis. During this process, the fusion of the late endosome with the lysosomal compartment is seen as a competing process to exocytosis, resulting in ILV degradation.^31^ In both processes, the canonical understanding of the lysosomal compartment is that of an endpoint for the degradation of biological material. However, recent work has highlighted the connection between the phagocytosis process and the MHC II antigen presentation mechanism.^14^ Phagocytosis by antigen-presenting cells often proceeds with the presentation of foreign antigen on cell surface MHC II receptors for the activation of downstream immune responses.^12^ This has led to the proposed link between phagosome fate and an MHC II-rich subclass of MVBs referred to as the antigen processing compartment.^14^ Consistent with this idea, prior infection studies have shown that professional phagocytes can secrete small EVs bearing MHC II loaded with bacterial cargo,^50^ suggesting a potential but incompletely defined link between phagocytosis and EV-mediated immune signaling.

Our work solidifies this connection, establishing a novel link between macrophage small EV secretion and particle phagocytosis. Across engineered cell lines, primary macrophages, and *in vivo* models, phagocytic update consistently amplified small EV secretion (**Figure 1**). Differences in the magnitude of the EV-increase in cell lines likely reflect metabolic differences and proliferation potential between cell types,^51^ which is known to influence EV output.^52^ This EV increase was independent of the macrophage phenotype (**Figure 6a**) and the dosed particulate chemistry (**Figure 2a**), suggesting that EV amplification is not a stress-induced response but unique to the phagocytic internalization route. However, particle size emerged as a critical determinant, with only particles large enough to undergo phagocytic uptake driving EV amplification (**Figure 2A**). Internalized particles localized within lysosomal compartments via TEM (**Figure 4**), consistent with canonical phagocytic trafficking and aligning with our previous work demonstrating that phagocytosis was the primary mechanism for PEGDA(-)50 internalization.^10^ Proteomic signatures in whole-cell PEGDA(-)50 dosed-BMM further supported phagocytic internalization and maturation, showing the increased presence of lynchpin phagosome proteins such as Arp2/3 complex-associated proteins, the WASH protein family,^53^ Cdc42,^54^ RILP,^55^ and SEPT2/11^56^ proteins.^34^ Together, this demonstration of phagocytic processing establishes a previously unreported upstream route for EV biogenesis, extending beyond traditional models that attribute EV formation solely to endosomal membrane trafficking or membrane budding.^1, 3, 4, 17^

Our data further indicate that phagocytosis-driven EV amplification primarily involves the biogenesis of small EVs originating from intracellular MVBs, aligning with current International Society of Extracellular Vesicles (ISEV) definitions.^1^ Through imaging (**Figures 3 and 4**), increases in MVBs colocalized with particle-containing lysosomes were consistently enriched following particle uptake, while increases of outward membrane budding indicative of microvesicle EVs were not observed across all particle types. Our gene enrichment analysis in PEGDA(-)50-dosed BMMs showed a persistent increase in proteins belonging to internal vesicle structures and organelles (**Figure 5E**), such as SNARE proteins, small GTPase, ESCRT-independent tetraspanin, and chaperone proteins,^3, 18, 31^ with an increased presence of proteins found in extracellular exosomes (FDR = 0.0358). Notably, in **Figure 4D ii.**, we see what appears to be the structure of the lysosome budding outward to a nearby MVB, suggesting transfer of lysosomal-degraded particulate products to the MVB for exocytosis. In line with enrichment data highlighting the importance of the lysosome and metabolic processing in these cells, our data suggests small EV loading can occur directly from phagolysosomes. While definitive assignment of EV subtypes and mechanistic pathways will require additional pathway-specific interrogation, these findings support a model in which phagocytic processing biases intracellular trafficking toward MVB formation and specifically small EV release. Future work should profile biogenesis markers found in phagocytosis-driven EVs, as well as incorporate inhibition studies, to develop the exact mechanism for phagocytosis-driven macrophage EV secretion.

Our use of the PEGDA NP platform is an enabling element of these studies. These hydrogel NPs are derived from PEG,^57^ a widely employed hydrophilic, biocompatible, FDA-approved polymer that does not elicit baseline immune activation in a variety of phagocytic cells, as demonstrated in prior publications,^10^ and strongly supported by our proteomic whole cell analysis (**Supplemental Figure S8** shows PEGDA(-)50 treatment compared to LPS inflammatory control). This lack of robust response to PEGDA NPs contrasts with other particle chemistries or bacterial probes, which have been shown to induce intrinsic immune activation,^10, 58^ leading to confounding effects between the known coupling of activation and phagocytic processing. While a growing body of literature has highlighted the generation of anti-PEG antibodies in animals and humans following exposure to PEGylated materials,^59^ such adaptive responses from multiple exposures are not expected to influence the acute exposures to naïve murine-derived cells presented in this work.

Using the synthetic control afforded by the PEGDA NP chemistry, we systematically decoupled particle number from total intracellular mass and lysosomal persistence. These studies revealed that small EV amplification depends on a critical mass of particulate material within macrophage lysosomes rather than the total number of internalization events. Particles with higher density (**Figure 2F**) or larger size (**Figure 2A**) induced comparable EV secretion when delivered at equal mass despite large differences in particle number, indicating that intracellular mass load is a primary determinant of EV biogenesis following phagocytosis. To further probe the intracellular persistence governing this effect, we employed acid-degradable PEGDA particles that undergo accelerated lysosomal degradation (*i.e.,* PEGDA(-)50/dS).^27^ Reduced lysosomal accumulation of degradable particles was accompanied by attenuated EV secretion, despite comparable initial uptake, demonstrating that prolonged residence of particle mass within the lysosome is required to sustain EV biogenesis (**Figure 2E**). All together from these studies, three critical particulate characteristics arise for the control of this phagocytosis-EV biogenesis mechanism: (i) a large enough particle size to facilitate internalization via phagocytosis (>200 nm)^43^, (ii) a minimal amount of particulate mass delivered to the cell (∼100 pg/cell), and (iii) an extended (> 24 hrs) residence time of accumulation in the lysosome.^27^

Beyond regulating EV quantity, phagocytosis-driven EV biogenesis also shapes EV composition and functional activity. EVs generated following particle uptake reflected the biological state of the source macrophages and could transfer this state to recipient cells, resulting in measurable changes in metabolic and immunological activity (**Figure 6**). Notably, recipient naïve macrophages exhibited phenotypic changes even when corresponding protein markers were not detectably elevated in the source EVs, suggesting that non-protein cargo contributes to EV-mediated signaling. Consistent with this possibility, EVs are known for also transporting lipids, RNA, and DNA across cells^17^ that can be selectively enriched relative to the cellular level, with distinct compositions between small and large EVs.^60, 61^ Furthermore, non-coding RNA in EVs has been shown to play significant roles in tissue health and disease progression.^62, 63^ Together, these findings indicate that phagocytosis-driven EV amplification enables the generation of functionally distinct EV populations that open new opportunities for therapeutic exploration. The potential to control EV composition in secreted EVs via particle amplification, either through modification of particle compositions to induce macrophage immunological response or to control metabolic processing and EV generation, would facilitate novel avenues for *in situ* drug delivery or *ex vivo* EV generation.^61, 64^ In particular, the increase in MHC II expression in secreted EVs highlights potential applications in antigen-dependent therapeutics, including vaccines, where enrichment of MHC II on EVs sourced from antigen-presenting cells has been shown to stimulate antigen-specific CD4+ T cells^65^ or facilitate transfer of antigen-loaded MHC II proteins through MHC II cross-dressing.^66^ More broadly, the ability to couple EV amplification with controlled cargo delivery established a novel avenue for macrophage-mediated EV therapies that leverage endogenous cellular communication pathways for targeted and sustained signal transfer.

In addition to transferring endogenous cellular components, phagocytosis-driven EV biogenesis enables the incorporation of exogenously delivered material into secreted EVs, expanding the functional potential of this pathway for both therapeutic design and EV manufacturing. Using fluorescently labeled particulate cargo, we demonstrate that material internalized through phagocytosis can be redistributed into distinct EV subpopulations and subsequently transferred to recipient macrophages (**Figure 7**). Notably, EV-mediated delivery produced distinct intracellular distribution patterns from direct nanoparticle uptake, as seen when comparing the Cy5 localization within the cell body in **Figures 7F** and **7H**. This difference points to fundamentally different internalization and processing routes for EV-mediated delivery as compared to particle-mediated. This distinct internalization method of EVs rather than particles could allow for unique drug delivery modalities, such as macrophage EV-mediated mRNA transfer, which has been shown to occur in both immediate and distal tissue sites.^67^ In contrast to earlier studies focused on sub-200 nm particles that enter cells via non-phagocytic routes,^67, 68^ our results identify phagocytosis as a productive and regulatable pathway for EV cargo loading that operates independently of macrophage activation state. Together, this mechanism establishes a framework for engineering macrophage-mediated EV delivery platforms that leverage endogenous phagocytic processing to achieve sustained, targeted transfer of therapeutic cargo to otherwise difficult-to-reach tissues. Future work should continue to investigate engineering particulate therapeutics that manipulate macrophage EV secretion for persistent release of cargo-loaded EVs and the potential to deliver to novel target tissues.

Through this work, we establish phagocytosis as a direct and previously unrecognized regulator of macrophage small EV secretion. EV amplification was governed by the accumulation and persistence of particulate mass within lysosomal compartments, suggesting that phagocytosis biases intracellular trafficking toward MVB formation and small EV release. This mechanism enables the incorporation of both endogenous and exogenous cargo into secreted EVs. Although additional studies will be required to fully resolve the molecular machinery underlying this process, the framework established here redefines the relationship between phagocytosis and EV biogenesis. Collectively, these findings explain fundamental phagocytosis processing and pave the path for a new design parameter that can leverage macrophage EVs in intercellular communication and therapeutic delivery.

## METHODS

### Cell Cultures

RAW264.7 murine cells and THP-1 human monocyte cells were purchased from American Type Culture Collection (ATCC, Manassas, Virginia, United States). RAW 264.7 cells were cultured between passages 2-10 in complete media, Dulbecco’s Modified Eagle Medium (DMEM) (Corning) supplemented with 10% heat-inactivated fetal bovine serum (FBS, Gibco), and 1% penicillin-streptomycin (P/S, Cytiva). THP-1 cells were cultured between passages 2-10 in complete media, RPMI 1640 Medium (ThermoFisher Scientific) supplemented with 10% heat-inactivated FBS and 1% P/S. Before experiments, THP-1 monocytes were terminally differentiated into macrophages by adding 200 nM Phorbol 12-myristate 13-acetate (PMA) for 24 hours, followed by a PBS wash and a 48 hrs resting period.

All studies involving animals were performed in accordance with National Institutes of Health (NIH) guidelines for the care and use of laboratory animals and approved by the Institutional Animal Care and Use Committee (IACUC) at the University of Delaware. Primary cells were extracted from female wild-type C57BL/6 mice (Jackson Laboratories) that were kept in a pathogen-free facility at the University of Delaware, with free access to chow. Previously reported standard protocols were used to generate BMMs isolated from mice.^69^ In brief, bone marrow was extracted from the femurs and tibias of euthanized mice and cultured in L929 fibroblast-conditioned media containing macrophage colony-stimulating factor.^70^ The collected bone marrow was strained, and cells were isolated using red blood cell lysis buffer. The bone marrow cells were then plated in 10 mL of differentiation media consisting of DMEM without L-glutamate and sodium pyruvate media, 20% FBS, 30% L929 conditioned media, and 1% P/S at 1*10^6^ cells/mL in a T-75 flask. On day 3, the cells were supplemented with an additional 5 mL of differentiation DMEM. Any excess bone marrow that was not plated on the day of collection (day 0) was then frozen down in freezing media, 90% FBS and 10% DMSO, according to previously reported protocol.^71^ On day 6, cells were ready for experiments. Experimental cultures used complete DMEM media (10% FBS and 1% P/S). Complete BMM differentiation was confirmed via CD11b, shown in **Supplemental Figure S13**.

### Polymeric Particles

PEGDA NPs were synthesized using a reverse emulsion technique previously described.^27^ In summary, solids PEGDA (Sigma Aldrich) (M_n_=700) (composition shown in **Table 1**), charge-establishing co-monomer (10 wt%), photoinitiator diphenyl (2,4,6-trimethylbenzoyl) phosphine oxide (TPO; Sigma Aldrich) (PI) (1 wt%), and fluorescent label sulfo-cyanine5 maleimide (Cy5, Fluoroprobes) (0.1 wt%) were dissolved in methanol (MeOH). For control of particle density, solid particle components were mixed at 50 wt% or 75 wt% with MeOH solvent. For negatively charged PEGDA particles, 2-carboxyethyl acrylate (CEA, Sigma Aldrich) was used as the co-monomer, while positively charged PEGDA particles used 2-aminoethyl methacrylate AEM (Sigma Aldrich). For PEGDA(-)50/dS, a thiol-PEG-thiol, M_N_=600 (Creative PEGWorks) was used as a co-monomer at 20 wt%, the detailed weight percentage in **Table 1**. We emulsified 100 μL of this monomer mixture in 1 mL of AP1000 silicone oil (Sigma Aldrich) and polymerized by irradiation with UV light (365 nm at ∼28 cm from the light source, ∼5-10mW cm^-2^) for 45 seconds. UV irradiation was performed using an APM LED UV Cube. Hexanes (Sigma Aldrich) were added to the polymerized mix to wash the silicone oil. Particles were then washed in ethanol at 16,000 rcf to separate unincorporated monomers and Cy5. Micrometer-sized particles were removed by passing the suspension through a series of sterile 5 μm and 0.45 μm filters. The resulting PEGDA NPs were then stored in ethanol at 4 ⁰C for future use. In experiments with non-PEGDA particles, 0.2 µm and 1 µm carboxylated PS particles (PolySciences) were obtained. Before dosing cells, particles were washed in sterile water prior to a final suspension in the appropriate experimental media.

### Particle Characterization

We performed thermogravimetric analysis (TGA) using TA Instruments TGA 550 to determine particle stock concentration following the final filtering step. We loaded 50 μL of particle suspension onto TGA sample pans in technical triplicates. The mass of the particle stock solution was determined via a mass reading after a temperature ramp to 90-120 ⁰C, followed by a 15 min isothermal step to ensure evaporation of ethanol or water solvent. Following determination of the stock concentration, dynamic light scattering (DLS) was performed using a Malvern Zetasizer Nano S Instrument (Malvern Instruments) to characterize the hydrodynamic diameters and zeta potentials of the particles. Preparation of DLS samples involved diluting to 0.1 mg/mL in deionized water. Hydrodynamic diameters and polydispersity indices were measured for technical triplicates.

### *In vitro* particle dosing studies

For characterization of dose dependent amplification of EV secretion, BMMs (1.5*10^6^ cells/well in 6-well plate, 3*10^6^ cells/flask in T-25, or 1*10^6^ cells/ 24-well plate), THP-1s (1.5*10^6^ cells/6-well), and RAW 264.7 (7.5*10^5^ cells/6-well) were seeded and allowed to adhere for at least 4 hrs. Cells were dosed at a range of particle concentrations and incubated with cells as outlined in figure captions. At designated timepoints, secreted EVs were isolated from culture media, while cells were processed for flow cytometry analysis of particle internalization and phenotype activation. LPS from Escherichia coli O111: B4 (Millipore Sigma) was dosed at 20 ng/mL as a positive control for BMM activation.

### EV isolation via differential centrifugation

Live cells were removed from the media by centrifugation at 180 rcf for 4 minutes. Cellular debris and apoptotic bodies were removed by centrifugation at 2,000 rcf for 10 minutes. Larger microparticles and remaining polymeric particles were removed by centrifugation at 16,000 rcf for 45 minutes. Following this last centrifugal step, the resulting supernatants were filtered through sterile 0.2 μm filters and centrifuged at 100,000 rcf for 90 minutes using a SW55 Ti rotor and Beckman Coulter ultracentrifuge L-90K or a TLA 55 rotor and Beckman Coulter Optima MAX Ultracentrifuge, depending on initial culture conditions. Pelleted EVs were resuspended in 1 mL PBS and washed once more using a TLA 55 rotor and Beckman Coulter Optima MAX Ultracentrifuge at 100,000 rcf for 90 minutes. Isolated EVs were then resuspended in 1 mL PBS and stored at 4 °C for further characterization and usage within a week. EV concentrations were measured using NTA on a NanoSight300.

### TEM of ultracentrifuge isolated EVs

For TEM analysis of ultracentrifuge isolated EVs, we used previously established negative staining method.^72^ Briefly, 400 mesh carbon coated copper grids were glow discharged with a PELCO easiGlow^TM^ glow discharge system to render the surface of the grids hydrophilic. Grids were briefly floated on drops of purified EV-PBS suspensions, washed on drops of water, and then negative stained with 2% uranyl acetate. Air-dried grids were examined with a Thermo Scientific Talos TEM operating at 120kV. Images were acquired with a Thermo Scientific Ceta 16M camera.

### *In vitro* BMM EV dosing

BMM cells were plated on 24-well plates (1*10^6^ cells/well) and given at least 4 hrs to adhere. After adhering, cells were dosed at 1*10^9^ particles/mL with ultracentrifuge isolated EVs from indicated source BMM culture conditions (EV^M0^, EV^M0-NP^, EV^M1-NP^, and EV^M2-NP^). After a 24 hrs incubation period, cells were processed for flow cytometry analysis of particle material transfer and phenotype activation.

### Flow cytometry analysis of *in vitro* cultures

Following outlined dosing conditions and incubation periods, macrophages were detached using 0.25% Trypsin-EDTA (Corning) and washed twice with FACS buffer (PBS supplemented with 2% FBS). Antibody staining of macrophages was performed following the initial two wash steps, otherwise samples were then analyzed using the ACEA NovoCyte Flow Cytometer. A complete table of antibodies used can be found in **Supplemental Table S1**. For surface antibody staining, BMMs were stained with I-A/I-E-Brillian Violet 785 (Biolegend) for 30 minutes on ice. After staining, samples were washed twice with FACS and analyzed using the ACEA NovoCyte Flow Cytometer. All PEGDA-based particle chemistries used contained a conjugated Cy5 molecule for measuring particle uptake and particulate material transfer through flow cytometry. Cells vs debris gate was determined by gating samples with high forward scatter area (FSC-A) and side scatter area (SSC-A). Singlets were isolated as having a 1:1 ratio of SSC-A and side-scatter height (SSC-H), as previously shown in literature.^73^ Untreated BMMs at each designated timepoint were used to set the NP+% gates and MHC II+%. Cy5 MFI was used to indicate the accumulation of NP material within source and naïve BMM cultures. Brillian Violet 785 fluorophore MFI was recorded via flow cytometry as a measure of MHC II expression in source and naïve BMM cultures.

### Magnetic bead Flow cytometry analysis of macrophage EVs

Following ultracentrifuge isolation of macrophage EVs, EV samples were processed for flow cytometry analysis, using the Basic Exo-Flow Capture Kit (System Bioscience, Palo Alto) as outlined in the vendor recommendation. Briefly, streptavidin-coated magnetic beads were conjugated with biotin anti-mouse capture antibodies in a 2 hrs incubation period. RAW 264.7 EVs were captured and analyzed using anti-mouse CD63, CD9, and ICAM-1 (**Supplemental Table S1**), while BMM EVs were captured and analyzed using anti-mouse CD63. Following attachment of biotin antibodies, isolated EVs were incubated overnight with the antibody-magnetic bead complex for sample captures. After overnight incubation, samples were washed twice and stained using the provided Exo-FITC or I-A/I-E Brilliant Violet 785 (Biolegend) for 2 hrs incubation, depending on the intended analysis. Samples were analyzed using the ACEA NovoCyte Flow Cytometer (Agilent Technologies). Cy5 MFI was recorded via flow cytometry as a measure of NP material transfer from source cells to secreted EVs. Brillian Violet 785 fluorophore MFI was recorded via flow cytometry as a measure of MHC II transfer from source cells to secreted EVs.

### Animal Studies

All studies involving animals were performed in accordance with NIH guidelines for the care and use of laboratory animals and approved by the IACUC at the University of Delaware. Female C57BL/6 mice of 2 to 4 months were housed in a pathogen-free facility at the University of Delaware and given unrestricted access to chow and water. Mice were dosed via orotracheal instillation following a standard procedure by suspension via their incisors and gently grabbing and moving the tongue aside to block the esophagus, allowing direct delivery to the trachea and minimal dosage to the gastrointestinal track. NP dosed mice were administered 100 µg PEGDA(-)50 in sterile 50 µL 1X PBS under anesthesia with isoflurane. AM uptake and EV BALF secretion analysis had a 72-hour endpoint. BALF was obtained via cannulation of the trachea restrained by a suture tie, using 1 mL of sterile 1X PBS dispensed into the lungs to withdraw and stored at 4 °C for further analysis.

After collecting, BALF was centrifuged at 500 rcf for 5 minutes. The resulting supernatant underwent differential centrifugation for further isolation of EVs (as above), while the pellet was resuspended in 500 µL red blood cell (RBC) lysis buffer (Fisher Scientific) and pipetted for 30-45 s. The suspended cells were then washed three times with 1X PBS at 500 rcf for mins. Primary antibody staining was applied to the samples, with antibody solution diluted in FACS buffer, and incubated for 45 mins on ice. Following the staining step, samples were washed twice with 200 µL of FACS before running with appropriate fluorophore channels continuously on the ACEA NovoCyte Flow Cytometer. Cells vs debris and singlet populations were determined as outlined in *in vitro* flow cytometry analysis (above). AM were identified as the Siglec-F^+^ population (BD Pharmingen).^73^ NP uptake was determined as NP^+^% based on the percentage of events with fluorescence intensity greater than 1% of the untreated population in the Cy5 channel.

### SEM of Particle-Dosed Macrophages

BMMs were seeded in ibiTreat Ibidi µ-Dish^35^^mm, high^ Grid-500 (Ibidi, München, Germany) at 1*10^6^ cells/dish, and allowed to adhere for at least 4 hrs. Samples were dosed with 100 µg/ml of various particle conditions: PEGDA(+)50, PEGDA(-)50, and PS(-)MP. After 24 hrs, samples were washed twice with 1X PBS and fixed using 2% EM grade glutaraldehyde (diluted from 8% Glutaraldehyde, Electron Microscopy Sciences) for a minimum of 1 hr, followed by three rinses in 1X PBS. Cells were dehydrated in an ethanol dilution series of increasing concentrations (25%, 50%, 75%, 95%, 100%, 100%). After dehydration, the cells were dried using an increasing dilution series of Ethanol: Hexamethyldisilazane (HMDS). Dilutions were as follows 2:1, 1:1, 1:2, 100% HMDS. Following the 100% HMDS step, the samples were allowed to air dry in a fume hood and then mounted to SEM stubs. Stubs were coated with 6 nm of platinum before being imaged at 2.0 kV on a Thermo Fisher Scientific Apreo Volumescope SEM.

### TEM of Particle-Dosed Macrophages

BMMs were seeded in Thermanox^TM^ Coverslips (ThermoFisher Scientific) at 3*10^5^ cells/dish and allowed to adhere for at least 4 hrs. Samples were dosed with 100 µg/ml of various particle conditions: PEGDA(+)50, PEGDA(-)50, and PS(-)MP. After 24 hours, samples were washed twice with 1X PBS and fixed using 2% EM grade glutaraldehyde, 2% EM grade paraformaldehyde in 0.1M sodium cacodylate buffer pH 7.4 (Electron Microscopy Sciences) for a minimum of 1 hr.

Following fixation, cells were washed 3 x 5 min with 0.1M sodium cacodylate buffer pH 7.4, post-fixed with 1% osmium tetroxide and 1.5% potassium ferrocyanide in 0.1M sodium cacodylate buffer pH 7.4 for 30 min, and then washed 4 x 5 min with water. The cells were *en bloc* stained with 1% uranyl acetate (aq) for 30 min, washed 3 x 5 min with water, and then dehydrated in an ethanol series (25% ethanol, 50% ethanol, 75% ethanol, 95% ethanol, and two exchanges in anhydrous ethanol, 5 min each step). Samples were gradually infiltrated with Embed-812 resin (25% resin in ethanol, 50% resin in ethanol, 75% resin in ethanol, and three exchanges in 100% resin) for 30 min each step. Coverslips were infiltrated in 100% Embed-812 resin overnight, and the next day were polymerized at 60 °C for 24h. The coverslips were separated from the polymerized resin by immersing them in liquid nitrogen. Areas of interest were excised using a jeweler’s saw and affixed to a flat-bottom BEEM capsule with cyanoacrylate glue. Cells were sectioned on a Leica UC7 ultramicrotome, and 60-70 nm thick sections were collected onto 200 mesh formvar/carbon-coated grids. The sections were post-stained with 1% uranyl acetate in 50% methanol and Reynolds lead citrate. Upon drying, grids were examined using a Thermo Scientific Talos TEM operating at 120kV. Images were acquired with a Thermo Scientific Ceta 16M camera.

### Proteomic and Mass Spectroscopy Preparation

BMMs were seeded in 24-well plates (3*10^5^ cells/well, 3 wells/repeat) and allowed to adhere for at least 4 hrs. Wells were dosed with PEGDA(-)50 at 100 µg/mL. After a 24 hrs incubation, cells were washed twice with PBS and incubated in incomplete DMEM for 48 hrs (no FBS). After this second incubation period, samples were collected, spun down at 500 rcf to remove cells, spun through an Amicon® Ultra-4 4 containing an Ultracel®-10 10K filter (10,000 NMWCO) at 5000 rcf to concentrate secreted proteins in 1 mL of incomplete DMEM media. A bovine serum albumin (BSA) protein assay (Bio-Rad) was then used to measure sample protein concentration before proteomics sample prep. Shotgun proteomics was completed according to a previously established method.^74^ Briefly, shotgun proteomics uses high-pressure liquid chromatography (HPLC) in tandem with mass spectrometry (LC-MS/MS) and database searching algorithms to identify proteins present in complex samples.^75^

BMMs were seeded in 24-well plates (3*10^5^ cells/well, 3 wells/repeat) and allowed to adhere for at least 4 hrs. Wells were dosed with PEGDA(-)50 at 100 µg/mL. Additionally, 20 ng/mL LPS was used as a positive control for macrophage cellular activation.^76^ After a 24 hrs incubation, cells were collected and prepped for in-cell proteomic processing. On filter, in-cell digestion was performed as described previously.^77^ Briefly, cells were fixed by methanol onto E4filters, followed by reduction, alkylation, trypsin protein digestion, and finally desalting. Desalted samples were dried and stored in -80 °C until LC-MS/MS analysis.^77^

### Proteomic Analysis

All generated raw files were submitted to Mass Spectrometry Interactive Virtual Environment (MassIVE, version1.3.16, University of California, San Diego). Upon obtaining proteomics analysis results, we used MaxQuant Pursuit 2.0.11 software for further data and statistical analysis. Of the total identified proteins, proteins that appeared in only 2 or fewer of the sample repeats were filtered out. Afterwards, imputations were added for any missing values within the remaining proteins. Values were determined using a normal distribution, with a width of 0.3 and a downshift of 1.8 around the identified proteins’ mean expression. Having filtered scarcely present proteins and filled in for missing sample values, a two-sample Student’s T-test was used to determine significantly different levels of protein expression, with a p-value ≤0.1 as the set threshold. Z-score values of the significant proteins were generated for future use in a hierarchical clustering heat map to identify upregulated protein clusters.

Volcano plots of the identified proteins were generated. Log fold change for each protein entry was calculated using the following equation:

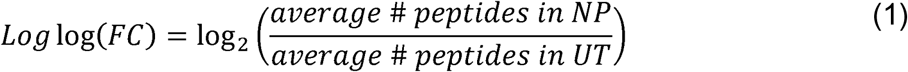

And significance was represented using:

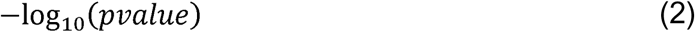

After identifying significantly different protein expression, biological network analysis was completed using Cytoscape_v3.10.0 stringApp.^78, 79^ Upregulated protein clusters were queried using their protein IDs, *Mus musculus* as their source, and edges represent confidence of connection greater than or equal to 0.4. Protein enrichment was completed using the provided stringApp tools and compared against genome wide background.

### Confocal Imaging of particle and EV-dosed macrophage cultures

BMMs were seeded in a 24-well plate (3*10^5^ cells/well) and allowed to adhere for at least 4 hours. After adherence, cells were dosed with 100 µg/mL PEGDA(-)50s or 1*10^9^ counts/mL EV^M0-NP^ and incubated for 24 hrs. Following incubation, BMMs were washed 2 times with PBS and fixed using 4% paraformaldehyde (PFA) before staining with Molecular Probe^TM^ Wheat Germ Agglutinin Conjugates (WGA) (ThermoFisher Scientific) according to manufacturer guidelines. WGA was used to stain the cellular membrane through the presence of N-acetylglucosamine and N-acetylneuraminic acid (sialic acid) residues. Briefly, wells were washed with PBS and then incubated in WGA working solution (5 μg/mL) for 10 minutes. Samples were then washed 2 times and stored in PBS until imaged on a Zeiss LSM 880 confocal microscope.

### Statistical Analysis

GraphPad Prism 9 (GraphPad Software Inc.) was used to perform statistical analysis, unless stated otherwise. All quantitative data are represented as mean ± standard deviation (SD). Data shown are from representative results from at least two independent experiments. Tukey’s or Dunnett’s multiple-comparison test, ordinary one-way ANOVA, or Student’s t-test were used to calculate p-values.

## ACKNOWLEDGEMENTS

The authors would like to acknowledge Yanbao Yu, Sylvain Le Marchand, Debbie Powell, and Shannon Modla for their assistance with proteomics and confocal imaging, respectively. We also thank the UD Office of Laboratory Animal Medicine (OLAM) for animal facility support.

## Author’s contribution CRediT statement

***M.T.R***.: conceptualization, data curation, formal analysis, investigation, methodology, software, validation, visualization, writing-original draft, writing-reviewing & editing; ***V.A.:*** investigation, validation, visualization; ***N.A.G.:*** investigation, methodology, validation; ***E.R.S.:*** investigation; ***C.A.F:*** conceptualization, funding acquisition, project administration, resources, supervision, writing-original draft, writing-reviewing & editing.

## Data and materials availability

All data needed to evaluate the conclusions in the paper are present in the paper and/or the Supplementary Materials. Proteomics data is archived at MassIVE submission MSV000100459. doi:10.25345/C56Q1SW81 and access link https://doi.org/doi:10.25345/C56Q1SW81. Figures 5G, 6A, and 6B were created with BioRender.

