## Supplemental Tables and Figures for "Particle Phagocytosis Amplifies Secretion of Small Extracellular Vesicles in Macrophages"

Author ORCIDs:

Trautmann-Rodriquez (0000-0002-2965-3537)

Sudduth (0000-0002-8141-6416)

Fromen (0000-0002-7528-0997)

*corresponding author

Catherine A. Fromen

150 Academy St.

Newark, DE 19716

(302) 831-3649

**Table of Contents:**


| | Page |
|---|---|
| **Supplemental Table S1:** Antibody table for flow cytometry experiments | S3 |
| **Supplemental Figure S1:** Example flow cytometry gating for RAW264.7, BMMs, and THP-1 | S4 |
| **Supplemental Figure S2:** Example flow cytometry gating for BALF isolated alveolar macrophage from *in vivo* mice studies | S4 |
| **Supplemental Figure S3:** Comparison between densities of PEGDA(-)50 and PEGDA(-)75 particle chemistries | S5 |
| **Supplemental Figure S4:** Second SEM imaging panel of BMMs dosed with distinct particle chemistry | S5 |
| **Supplemental Figure S5:** Second TEM imaging panel of BMMs dosed with distinct particle chemistry | S6 |
| **Supplemental Figure S6:** Image analysis of vesicle per cell in TEM imaging of particle-dosed BMMs | S6 |
| **Supplemental Figure S7:** Volcano plots for whole cell and secreted proteomic analysis | S7 |
| **Supplemental Figure S8:** Hierarchical clustering heat map for significantly differentiated proteins in UT, PEGDA(-)50, and LPS-dosed BMMs whole cell proteomic analysis | S7 |
| **Supplemental Figure S9:** Metabolic activity of PEGDA(-)50 and $EV^{MO-NP}$ dosed BMMs using CellTiter Glo 2.0 | S8 |
| **Supplemental Figure S10:** Confirmation of pro-inflammatory BMM phenotype after 72 hrs incubation | S9 |
| **Supplemental Figure S11:** Confirmation of anti-inflammatory BMM phenotype after 72 hrs incubation | S10 |
| **Supplemental Figure S12:** Example flow gating of Cy5 expression in phagocytosis-driven BMM EVs after 72 hrs of incubation | S11 |
| **Supplemental Figure S13:** Example flow cytometry gating for BMMs and CD11b marker | S11 |

**Supplemental Table S1:** Antibodies for flow cytometry analysis. Antibodies for staining and magnetic bead capture were purchased through Biolegend with corresponding fluorophore, clone number, and dilution factor listed.

| Marker of Interest | Distributor | Fluorophore | Clone | Dilution |
|---|---|---|---|---|
| CD11b | Biolegend | Pacific Blue | M1/70 | 1:100 |
| CD80 | Biolegend | Pacific Blue | 16-10A1 | 1:100 |
| MHC II | Biolegend | BV 785 | M5/114.15.2 | 1:100 |
| CD163 | Biolegend | BV 421 | S15049I | 1:100 |
| Siglec-F | BD Pharmingen | APC-Cy7 | E50-2440 | 1:100 |
| CD63 | Biolegend | NA | NVG-2 | NA |
| CD9 | Biolegend | NA | MZ3 | NA |
| ICAM-1 | Biolegend | NA | YN1/1.7.4 | NA |

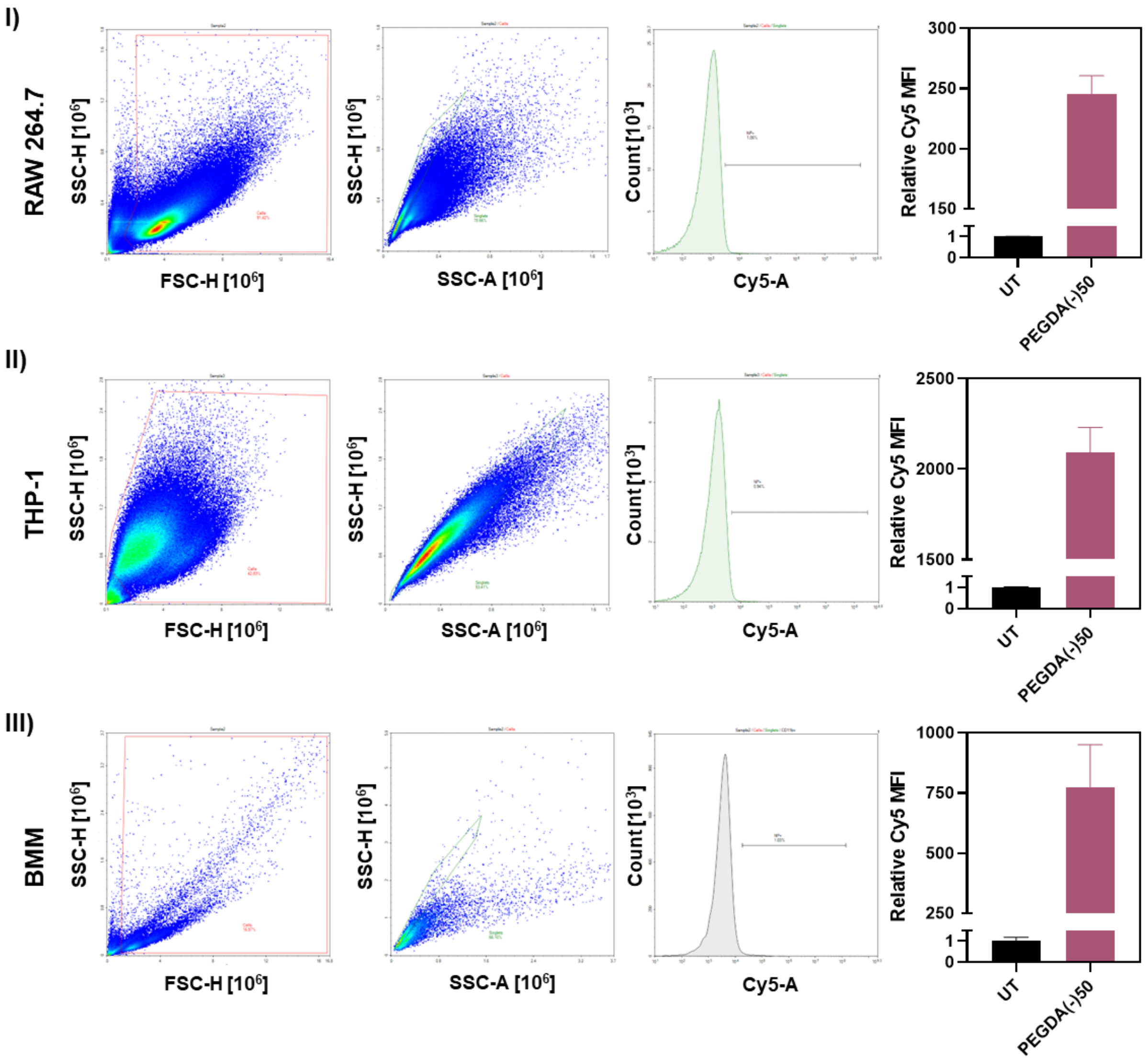


**Supplemental Figure S1** Example flow gating (I) for RAW264.7, (II)THP-1, and (III) BMM cultures with Cy5 MFI for untreated and PEGDA(-)50 dosed with (100 µg/mL) cultures.

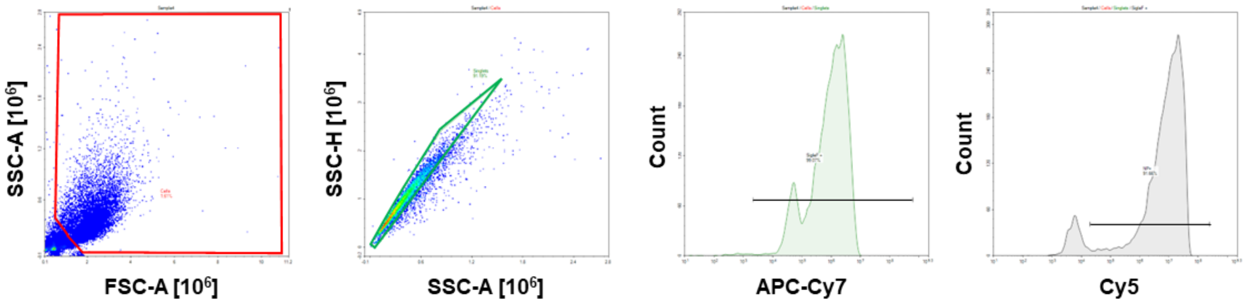


**Supplemental Figure S2.** Example flow gating for alveolar macrophages isolated from PEGDA(-)50-dosed mice. Density plots selecting for cells, singlets, and Siglec-F+, and showing PEGDA(-)50 internalization.

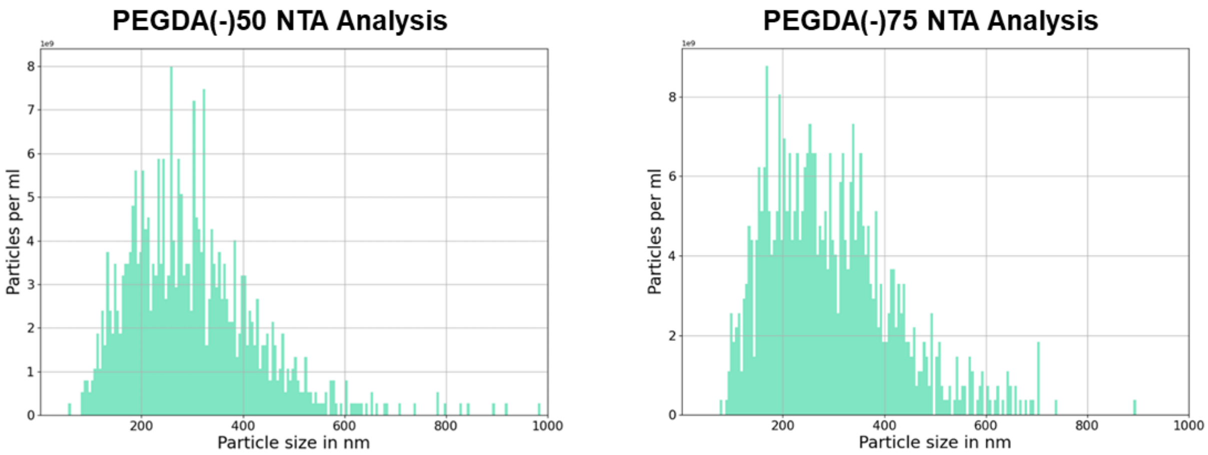


| Particle Chemistry | Diameter (nm) | Particle count (particle/mL) | TGA Mass (µg) | Volume ($nm^3$) | PEGDA Mass /particle (mg/particle) | Particle PEGDA Density (kg/m^3) |
|---|---|---|---|---|---|---|
| PEGDA(-)50 | 283.3 | 2.59e11 | 52 | 9.52e7 | 4.02e-12 | 42.16 |
| PEGDA(-)75 | 277.4 | 3.93e11 | 155 | 8.94e7 | 7.86e-12 | 87.88 |

**Supplemental Figure S3.** NTA and TGA of PEGDA(-)50 and PEGDA(-)75 chemistry, used to calculate particle density based on diameter, stock particle counts, and stock concentration. Volume assumed to be spherical.

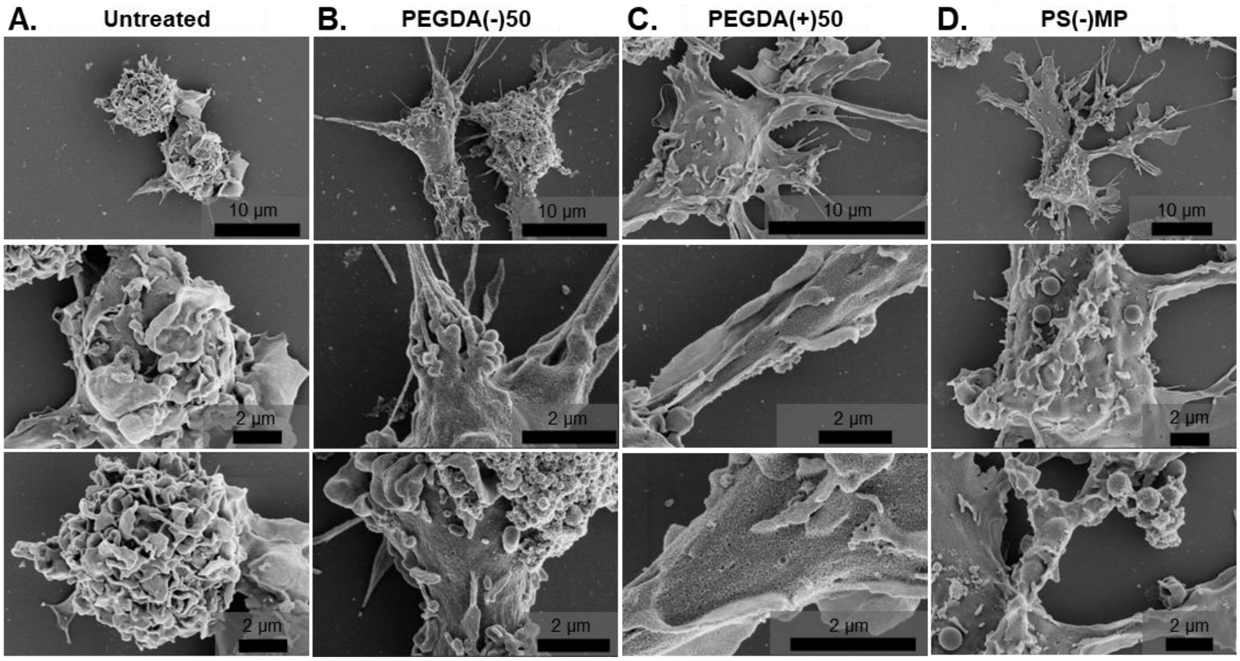


**Supplemental Figure S4.** Replicate set of SEM images for **A)** untreated, **B)** PEGDA(-)50, **C)** PEGDA(+)50, and **D)** PS(-)MP BMM cultures.

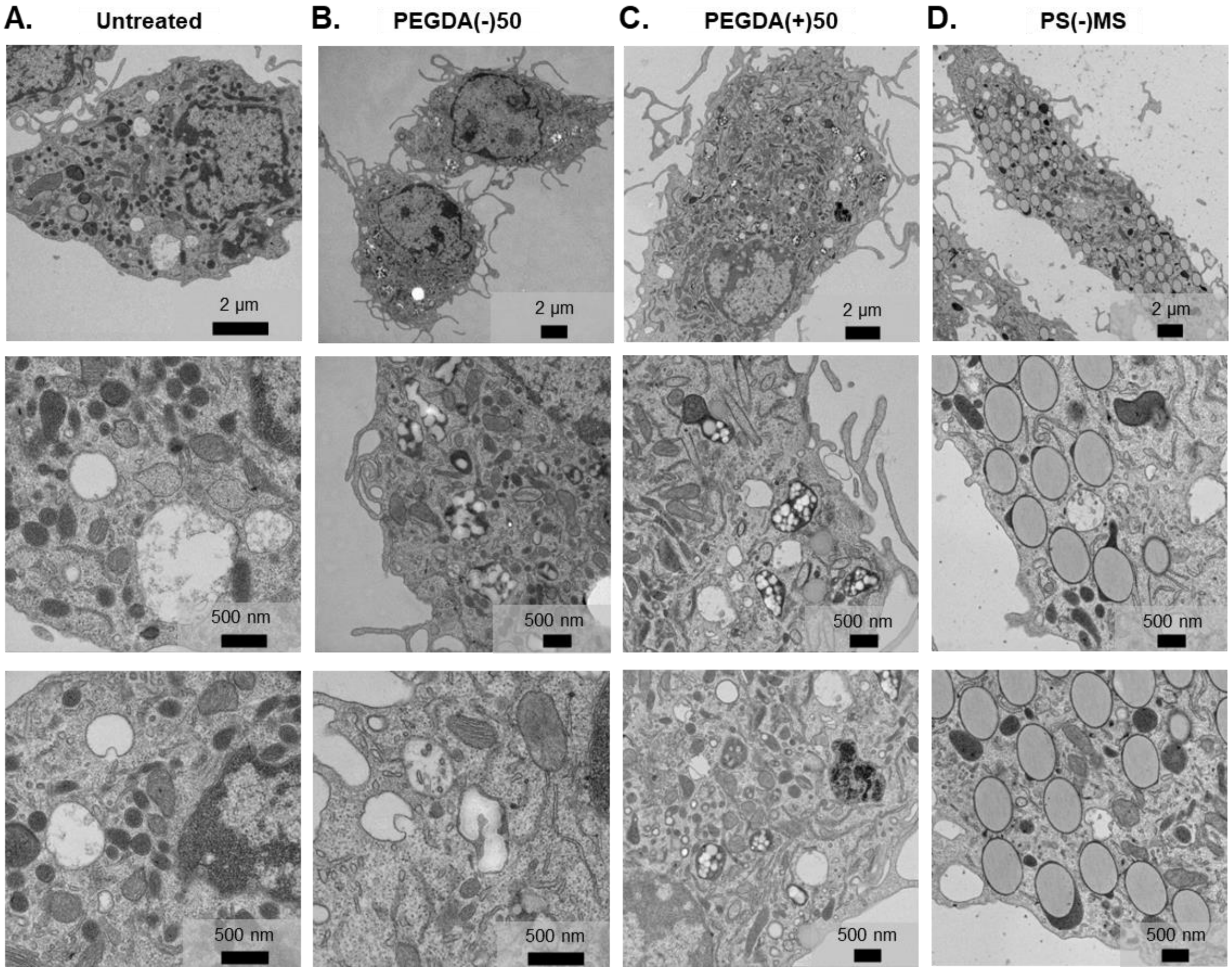


**Supplemental Figure S5.** Replicate set of TEM images for **A)** untreated, **B)** PEGDA(-)50, **C)** PEGDA(+)50, and **D)** PS(-)MP BMM cultures.

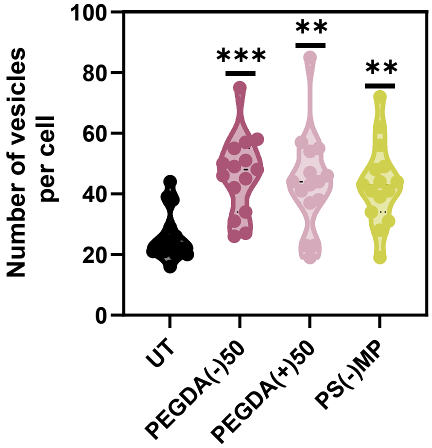


**Supplemental Figure S6.** Graph showing the number of vesicles in each cell, as calculated from the TEM images, PEGDA(-)50(p-value) = 0.0005, PEGDA(+)50(p-value) = 0.0025, and PS(-)MP(p-value) = 0.0031 (ordinary one-way ANOVA with a Turkey's multiple comparison test, N = 15).

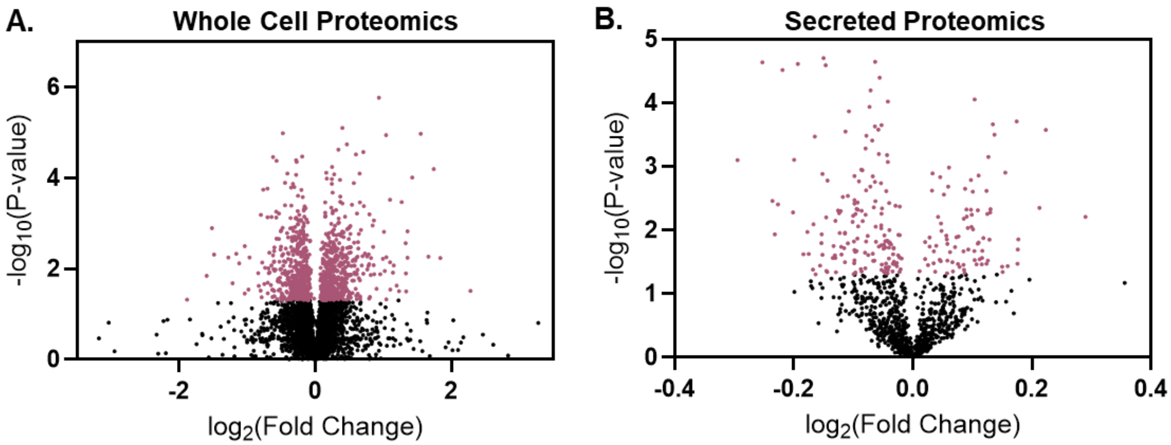


**Supplemental Figure S7.** Volcano plots for identified proteins in **A)** whole cell proteomic study and **B)** secreted proteomics study.

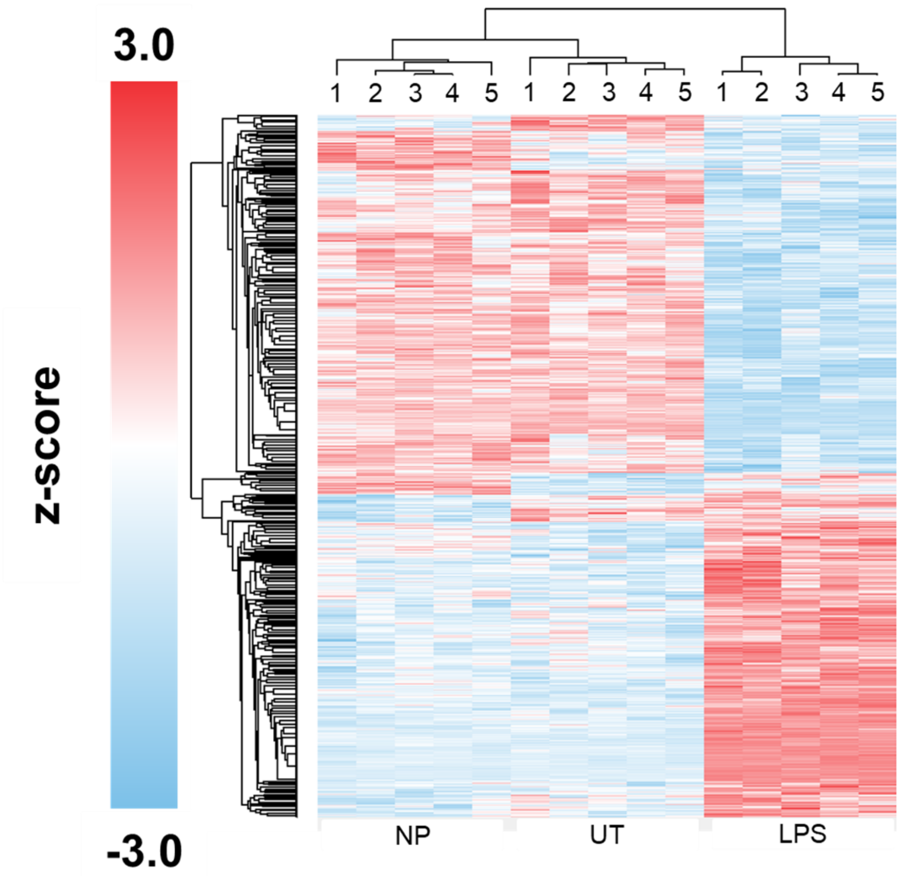


**Supplemental Figure S8.** Hierarchical clustering of proteomic datasets comparing protein expression in UT, PEGDA(-)50-dosed, and LPS-dosed BMMs. Clustering highlights similarity between UT and NP, when compared to inflammatory (LPS-dosed) BMMs.

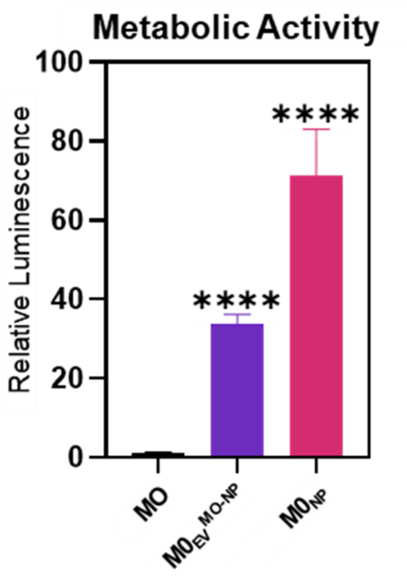


**Supplemental Figure S9.** Metabolic activity of BMM cultures after 4 days of dosing with PEGDA PEGDA(-)50 and NP derived EVs as measured through Cell Titer glo 2.0 (N = 4, SD error bars) ****p<0.0001, using Brown-Forsythe and Welch ANOVA tests.

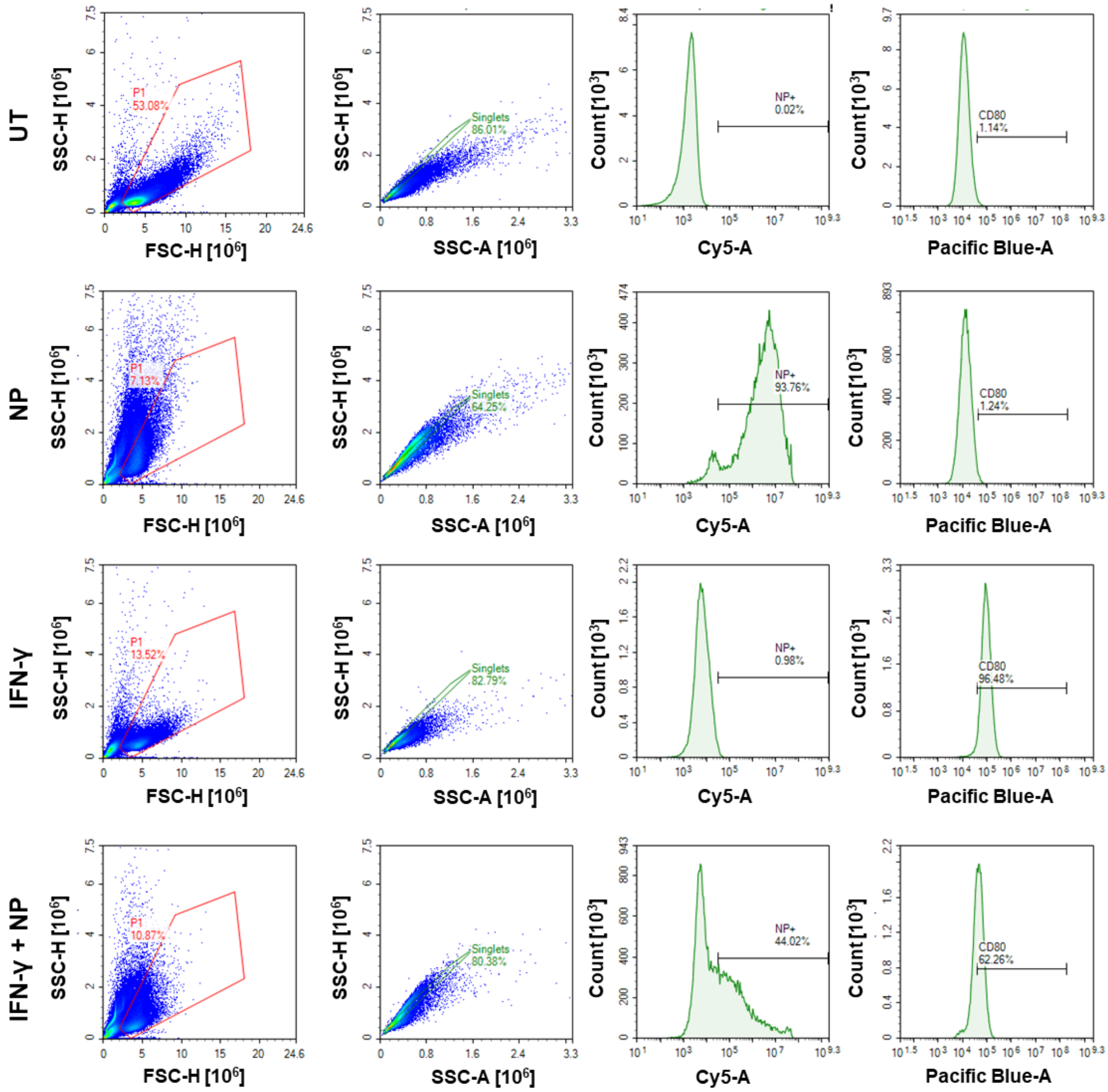


**Supplemental Figure S10.** Example flow cytometry plots of BMM samples following dosing and 72 hrs incubation with PEGDA(-)50, IFN-γ, or PEGDA(-)50 and IFN-γ. Pro-inflammatory marker CD80 highlighted here.

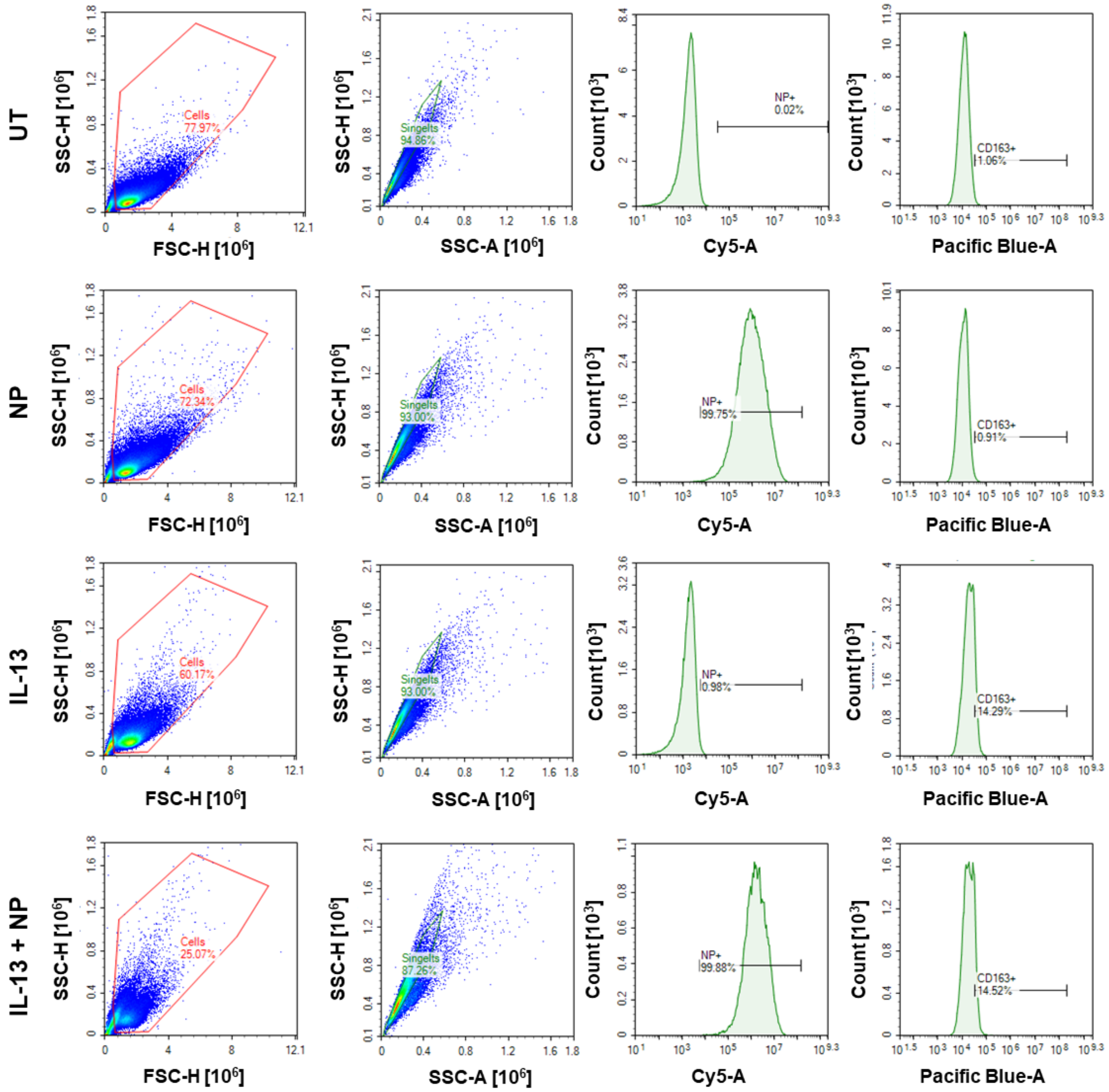


**Supplemental Figure S11.** Example flow cytometry plots of BMM samples following dosing and 72 hrs incubation with PEGDA(-)50, IL-13, or PEGDA(-)50 and IL-13. Anti-Inflammatory marker CD163 highlighted here.

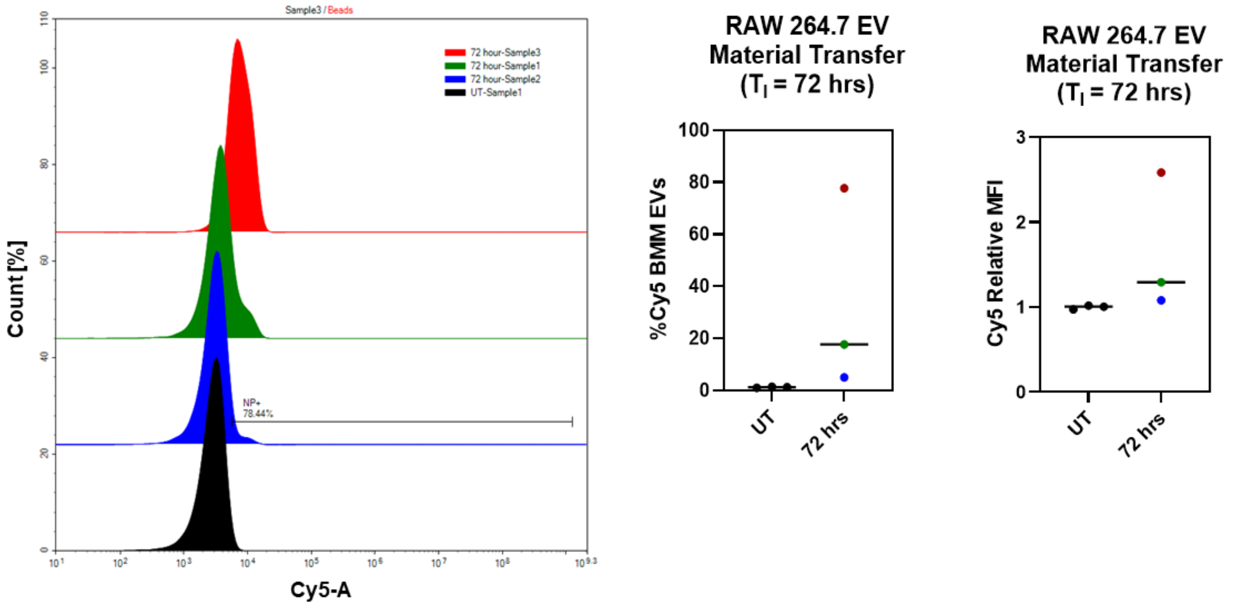


**Supplemental Figure S12.** Anti-CD63 capture bead flow cytometry analysis, sample histograms, relative Cy5 MFI, and %Cy5+ BMM EVs, in PEGDA(-)50 amplified EVs as a marker for NP material transfer for $T_I$ = 72 hrs.

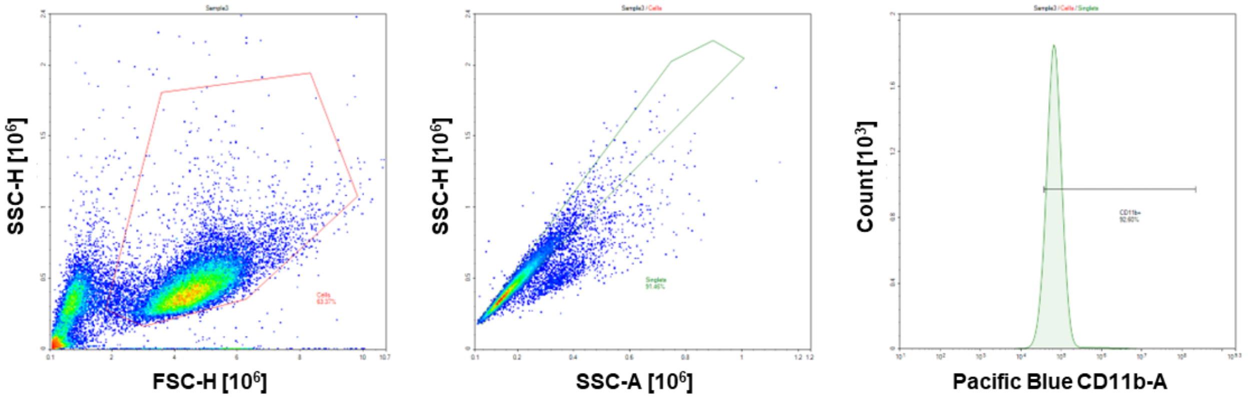


**Supplemental Figure S13.** Example flow gating for BMMs, with BMM marker CD11b+ highlighted.
